# Distortion Discovery: A Framework to Model, Spot and Explain Tumor Heterogeneity and Mitigate its Negative Impact on Cancer Risk Assessment

**DOI:** 10.1101/2021.04.28.441787

**Authors:** Dalia Elmansy

## Abstract

In a complex system of inter-genome interactions, false negatives remain an overwhelming problem when using omics data for disease risk prediction. This is especially clear when dealing with complex diseases like cancer in which the infiltration of stromal and immune cells into the tumor tissue can affect the degree of its tumor purity and hence its cancer signal. Previous work was done to estimate the degree of cancer purity in a tissue. In this work, we introduce a data and biomarker selection independent, information theoretic, approach to tackle this problem. We model distortion as a source of false negatives and introduce a mechanism to detect and remove its impact on the accuracy of disease risk prediction.

## 1. INTRODUCTION

Data that does not convey meaning and cannot be interpreted correctly by machines is considered noise. Sources of data noise are myriad. Data noise results in a range of problems from waste of space to skewness of reality and inaccuracy of predictive results. Some of the known sources of data noise include data collection, entry, transmission or inconsistencies of used conventions (e.g. naming, coding, format, etc.) ^1^. Accordingly, there are many different ways of cleaning up data for purposeful use. Such ways of handling noise in data depend on many factors including the data source, type, sensitivity and purpose.

Noise in omics data is particularly complex. From one side, this is due to the multitude of its sources, many of which are unknown and hence the noise remains unexplained. The complexity, from the other side, is due to the adverse effects of data noise on the accuracy of disease risk predictions.

Lazar et al. ^3^ differentiate between three types of noise in omics data: heterogeneity, batch effect and bias. Heterogeneity is due to the effect of other unobserved technical components of the study that lead to “variability of the outcome”. They differentiate it from batch effect, which is also technical, but more systematic. The latter occurs when data is prepared and used in different lots, is also unrelated to any biological variation under study, but rather, it could be due to, sampling or processing time, method of data collection, hybridization ^3^. Bias on the other hand, includes other distorting factors that accidentally distort the association between the investigated factors and the biological variation of interest ^2^. Integration of multi-level omics data, while needed, introduces various potential experiment-related sources of data noise.

Due to the complexity of biological systems and the intricacy of inter-genomic interactions, the lack of samples and the high degree of noise in omics data, false predictions remain an over-powering problem when using omics data for disease risk prediction. This problem is especially clear when dealing with complex diseases like cancer.

In cancer, the purity of the tumor sample is sometimes undermined as a result of the contamination of the tumor tissue with other non-tumor cells. Such impurities in the tumor tissue, for e.g. normal epithelial, connective tissue cells, immune cells and vascular cells lower the tumor content of the tissue, which in turn interferes with the sample’s tumor signal ^4^ and its usefulness for risk assessment. Typically, machine learning algorithms would drop such samples in the training phase, considering them as unexplained noise. If there is a significant number of impure samples, as well as insufficient array of input samples, this could interfere with the predictive ability of the classifier, and especially results in a large number of false negatives predictions.

Some methods have been previously suggested to estimate the purity of cancer content in tumor tissues either by using DNA copy number data, e.g. ABSOLUTE ^5^, or by using next-generation sequencing data, e.g. PurityEst ^6^. DNA copy number-based estimation of tumor purity is very popular for its accuracy, however, it works on copy number data which makes it limited to its availability. Some other methods sort out the gene expression data into profiles based on its cellular components. Other methods exploit the differences in transcription behavior of evident cell types. Such methods create a profile for each evident cell type in a normal tissue, taken from microarray data, by calculating an enrichment score for each cell type^7–11^. An example of the latter is ESTIMATE (Estimation of STromal and Immune cells in Malignant Tumor tissues using Expression data)^4^, which is another popular method. ESTIMATE uses gene expression data of tumor samples, and employing knowledge of cancer distinctive transcriptional gears, they focus on two types of cells that they presume make up the majority of impurities in the tumor sample; the stromal and immune cell types. They define two profiles, one for stromal and another for immune cells. They calculate a score for each profile using single sample gene set enrichment analysis from which they deduce tumor cell purity.

Other work was done to estimate the degree of cancer purity in a tissue using gene methylation data. These include LUMP (leukocytes un-methylation for purity) ^12^, InfiniumPurify ^13, 14^, PAMES (Purity Assessment from clonal MEthylation Sites)^15^ and RF_Purify ^16^. LUMP calculates the mean values of the CpG sites that are well known to be hypo-methylated in immune cells. InfiniumPurify uses kernel density estimation of the tumor purity after figuring out the differentially methylated CpG sites by analogizing tumor and normal samples. PAMES works on CpG islands, they compute the average methylation value for every CpG island, then the area under the Receiver Operating Characteristic (ROC) curve for each CpG island based on which they decide if a CpG site is to be included in the model or not. They estimate the tumor purity from the median of hypo-methylated and hyper-methylated sites. RF_Purify estimates tumor purity without using a reference (a set of control samples) by training two random forest classifiers on ABSOLUTE ^5^ and ESTIMATE ^4^ tumor purity scores of samples taken from The Cancer Genome Atlas (TCGA) ^17^.

In this work, we introduce a dataset and biomarker selection-independent framework to tackle this problem of tumor tissue impurity and to mitigate its negative effect on prediction, especially the false negative rate. We model distortion as a source of false negatives and introduce a mechanism to discover it, explain it and remove its impact on the accuracy of disease risk prediction. We apply the model to prostate cancer, as a test case, and we use multiple gene expression datasets (Prostate Cancer case/control, subtypes, metastatic/non-metastatic, aggressive/non-aggressive) of different platforms (microarray expression, RNA-Sequencing, DNA methylation) in order to interpret the hidden factors underlying the misclassifications and hence improve classification accuracy. We show that the prediction accuracy of the introduced model is better than some of the state of the art machine learning algorithms, in a simple, scalable, and interpretable fashion^18^.

**Figure 1** shows a high level block diagram of the distortion discovery algorithm and its validation.

**Figure 1.**
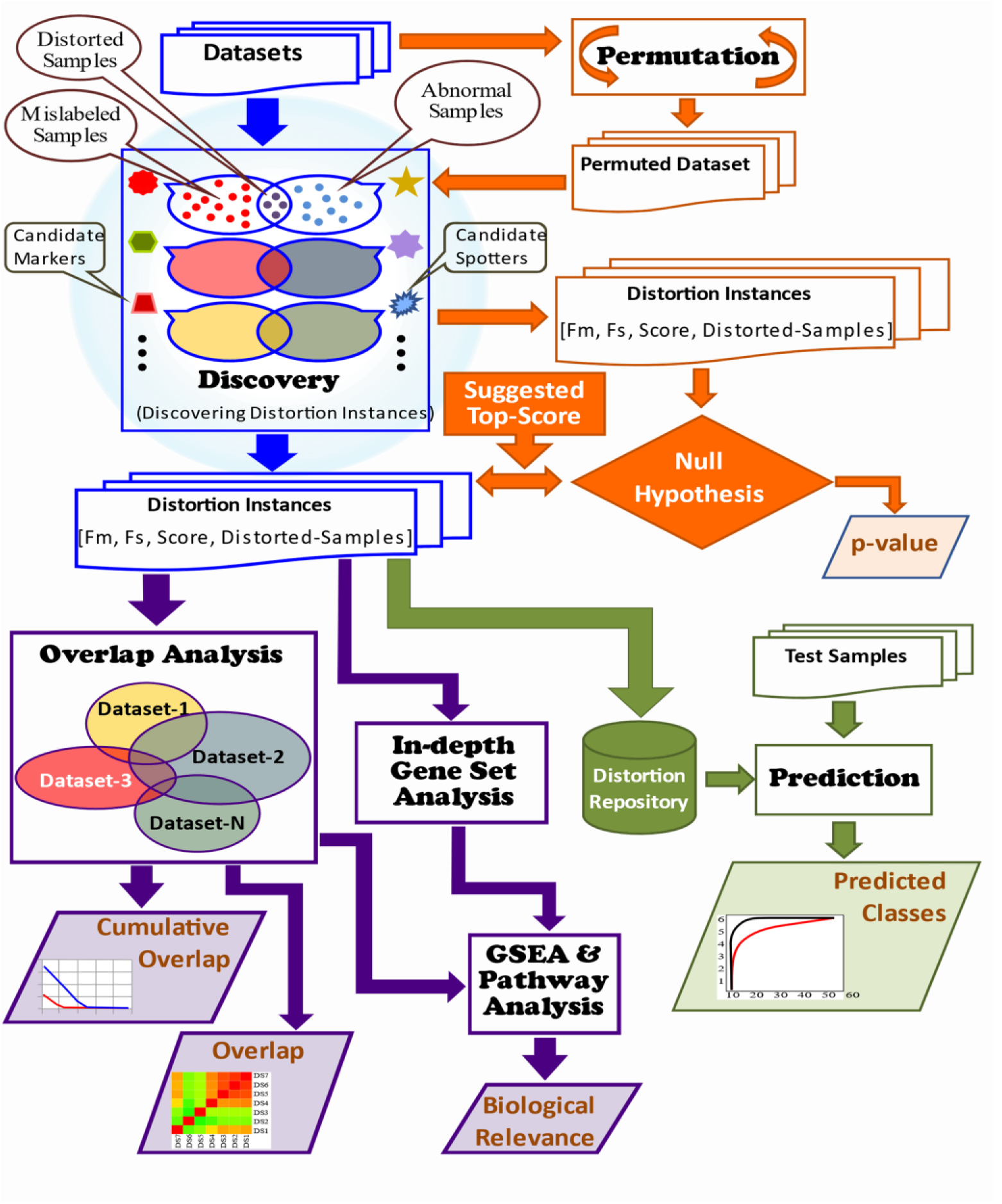
High Level Block Diagram of Distortion Discovery and Correction. The figure shows four pipelines: Discovery (in blue): discovers distortion instances and corrects for distortion using the proposed method. The three other pipelines are assessments: of significance (Permutations in orange), utility (Prediction in green) and biological relevance (Overlap Analysis in indigo).

**Figure 2.**
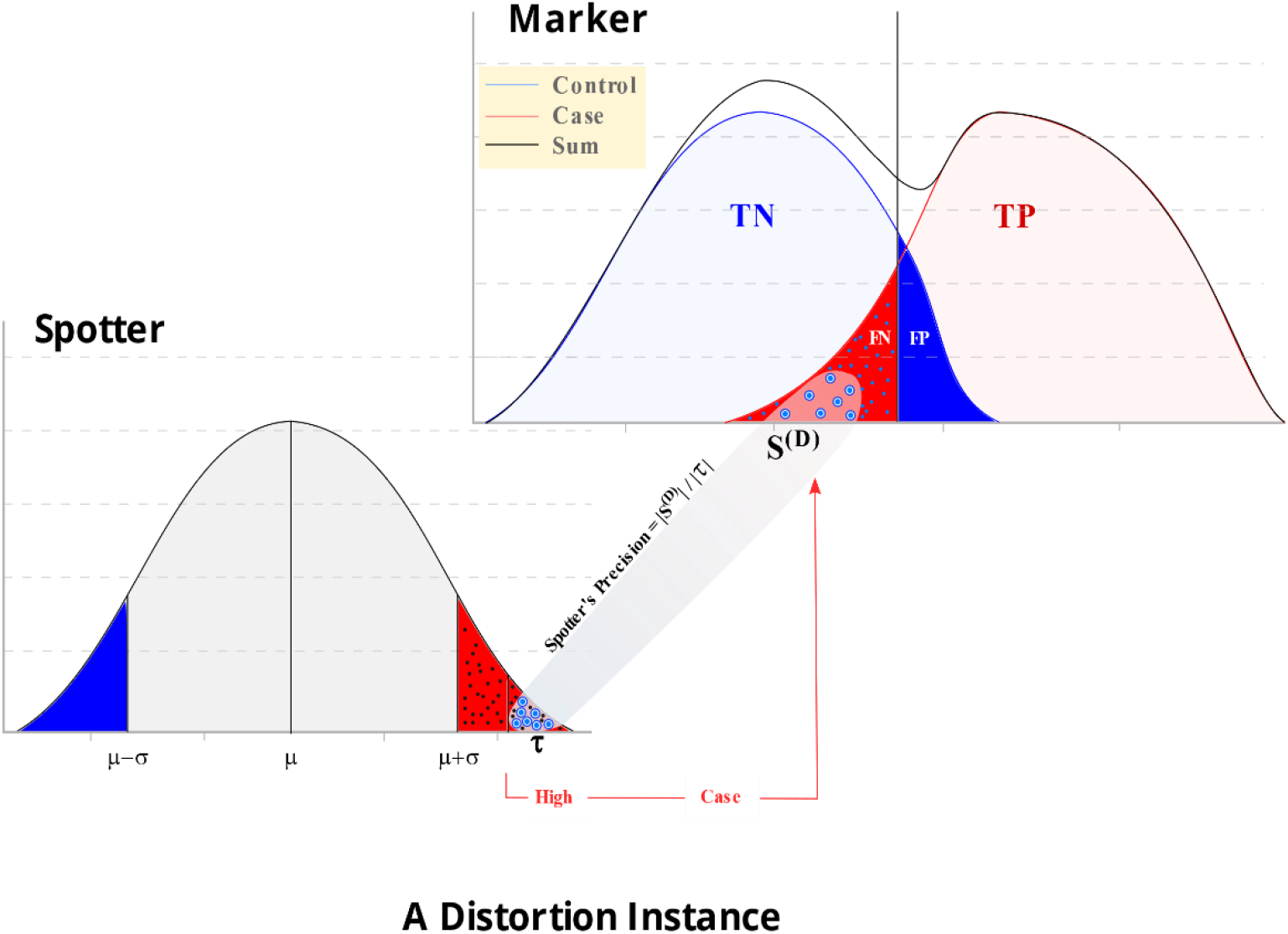
A Distortion Instance. The figure shows the distributions of the expression values of a marker gene and a spotter gene in case and control samples. The spotter gene is spotting a subset of the false negative samples in the distorted marker’s context with a spotter’s precision = 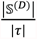, representing the fraction of the marker’s distorted samples in the spotter’s high tail.

## 4.2 METHODS

The objective of this study is to model, discover, correct for and try to characterize factors that are potentially associated with noise or bias inherent to omics data. We model distortion–a portion of noise - in the identification of biomarkers and introduce a mechanism to characterize it, discover it and correct for its effect. In order to do that, we define candidate distorted markers, candidate distorted spotters and distortion instances:

- A candidate distorted marker is a factor, usually molecular markers identified using omics data, with compromised predictive ability, which could be due to the effect of distortion.
- A candidate spotter is a factor used to spot the (potentially systematic) effect of distortion on one or more candidate distorted marker(s).
- A distortion instance is a set consisting of a distorted marker, its distorted samples, the actual class of distorted samples, the spotter that spots the distortion, and the spotter’s category (e.g. up-regulated or down-regulated in the phenotype of interest) as well as the spotter’s precision in spotting its coupled marker’s distortion in the dataset.

In this section, we first model the relationship between a marker and its spotters. We then design an algorithm that discovers and scores distortion instances in a dataset and develops predictive models by correcting for the effect of distortion. In this work, we use single gene markers that are defined using information theory. The introduced criteria for correction is independent of any underlying dataset or the criteria used to identify markers. Once we identify distortion instances, we systematically evaluate their utility in improving the predictive ability of markers and characterize the functional commonalities and relationships between spotters discovered in different datasets.

### 4.2.1 Defining distorted markers

A **marker gene** is a gene whose behavior (e.g., mRNA-level expression) is associated with the studied phenotype. If, for example, the phenotype in question can take two categorical values (referred to as “class” throughout this chapter), the expression of a marker gene should ideally have two distribution curves that are clearly separable from each other, potentially intersecting at the tails (**Figure**).

The extent of overlap between the tails of the two curves depends on how differently the marker gene behaves in each class. The more the overlap, the less differentiated the marker’s behavior in the two classes. So if the marker gene is used for classification, i.e. to predict the class of a given sample, then it would be less accurate as the overlap between the tails increases. On the other hand, the marker would be more accurate in predicting the class of a sample as the overlap between the tails decreases, which means less number of misclassifications, i.e. sample mislabeling.

In the following discussion, for the sake of simplicity, we focus on single gene markers. However, recognizing the limitations of individual genes in serving as markers, particularly for predictive tasks, the proposed method can be applied to markers composed of pairs of genes as well as sets of multiple genes.

### 4.2.2 Identifying candidate markers using information theory

#### Definitions

**Entropy** is a measure of uncertainty. In information theory ^19, 20, 45^, the uncertainty of any random variable X (the random variable could be a genetic variant, a gene expression, a protein, an epigenetic factor, etc.) that can take either discrete or continuous values, is measured by its entropy ***H*(*X*)** as follows:

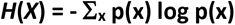

If the random variable is discrete, for example binary, the entropy would be calculated from the probabilistic model of the relative frequencies for each value of x.

**Conditional entropy** is the uncertainty of a random variable *X* given knowledge of another random variable *Y* and is denoted by ***H*(*XIY*)** as follows:

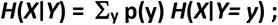

where *H*(*X*|*Y= y*) = - Σy p(x|y) log p(x|**y**)

**Joint entropy** is the uncertainty of a pair of random variables *X*, *Y* denoted by

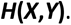

The relationship between the above three quantities is shown in the equations below:

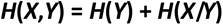

**Mutual Information** is a measure of dependency between two variables. It is the information gain or the reduction of uncertainty by/of one variable as a result of the knowledge of the other. So for two random variables *X* and *Y*, the amount of information that each one of them provides about the other is the mutual information ***I*(*X*;*Y*)**. For data drawn from the distribution p(X,Y), ***I*(*X*;*Y*)** quantifies the expected log-likelihood ratio of the joint probability p(X,Y) as opposed to the product of their individual probabilities p(X)p(Y). It is calculated from entropy and conditional entropy as follows:

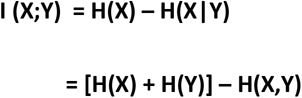

The mutual information is intuitively a non-negative quantity. It is zero if and only if *X* and *Y* are independent of each other. If the variables are normalized, mutual information takes values from zero to one.

It is important to note that if the variables are continuous, then the mutual information needs to be estimated. This requires an estimate of the probability distribution underlying the data, which should to done carefully such that it does not bias the resulting mutual information value, especially in small sized datasets. Many applications have used practical estimation techniques of mutual information on continuous data ^21–26^ a comparison of which showed that kernel density estimators and k-nearest neighbors (for k=3) performed best on smaller datasets with noise ^21^.

**Synergy** is defined by a cooperative creation of a whole that is greater than the sum of its parts. So the synergy of two random variables X and Y with respect to an outcome Z is the difference between the collective effect of the two random variables together and the sum of each of their individual effects. Synergy can be defined from mutual information as follows ^45–27^.

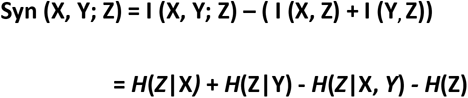

The synergy equation above quantifies the effect on the outcome variable Z due to the cooperative effect of the two random variables X and Y. A positive synergy means that there is an advantage to using X and Y together in identifying the effect on the outcome Z over using the sum of each of the variables’ effect alone. A zero synergy means no cooperation between the two variables in identifying the effect on the outcome. A negative synergy implies that each of the two random variables X and Y can replace the other in getting the same total effect on the outcome variable Z, i.e. redundancy. The synergy formula can be generalized to a multivariate formula and a tree of synergies representing the hierarchical relationships between a set of synergistic subsets of variables ^45–47^.

In this work, we use the definitions of entropy and conditional entropy above to compute the mutual information of continuous gene expression values and the phenotype of interest. We also compute the synergy between our distorted markers and their spotters with respect to the phenotype of interest, in order to explore if there is any biological association between the distorted markers and their spotters.

#### Benefits of using information theoretical measures

In this work, we use mutual information to identify individual candidate markers because we believe it is an intuitively simple and powerful tool, and makes good sense of large data. Mutual information, as a measure of statistical dependency, is capable of capturing linear and non-linear relationships. It is symmetric and does not make any assumptions about the distribution of the variables. Mutual information is also robust to outliers, can be generalized to more than two variables and its self-equitability characteristic as defined by ^27^ makes it able to quantify associations without bias for relationships of one type or another and it has been empirically proven to have a higher statistical power as opposed to other measures of association ^28^.

#### Previous Work

Anasstassiou and Varadan ^45–47^ previously used mutual information to gain insights into the information that the expression level of a gene *G_i_* in a tissue provides about the presence of a phenotype of interest *C* given datasets in both presence and absence of the phenotype. They quantified the information that any gene provides about the phenotype using mutual information as follows:

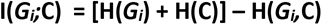

Subsequently, they defined the information **I(*G_i_*,*G_j_* ; *C*)** that any pair of two genes (*G_i_*, *G_j_*) jointly provide about the phenotype of interest *C*, as well as the information that ***n*** genes I(*G_i_*,*G_j_,.. Gn* ; *C*) jointly provide about the phenotype of interest *C*. Hence genes can be prioritized (or pair of genes or sets of multiple genes) with respect to the phenotype of interest. They observed that it is common that high-ranked gene sets do not include any of the high-ranked single genes, which indicated that the association of the genes with the phenotype, is due to a “*purely cooperative”* effect of the genes ^46, 47^. They quantified such cooperative effect using synergy which they defined for a pair of genes with respect to a phenotype C as:

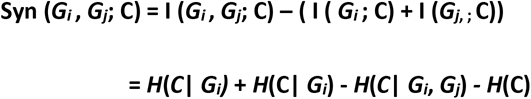

Watkinson et. al ^48^ used gene pairs as markers, and calculated synergy of each pair of genes. For this purpose, they used the unweighted pair group method with arithmetic mean (UPGMA) clustering algorithm ^29^ to estimate mutual information from the genes’ joint expression levels data in the presence and absence of prostate cancer, after normalizing the data using the Robust Multi-array Average (RMA) method ^30^ on perfect match probes. In order to calculate synergy of each pair of genes, they clustered the samples in the Cartesian space of the gene pair, then they used an empirical cutoff to define the samples’ clusters. For each cluster they calculated the entropy of the cluster as: *h*(*Q*) = -*Qlog*_2_ *Q* - (1-*Q*) *log*_2_(1-*Q*), where *Q* is the relative frequency of cancerous samples in the cluster. Then the entropy of the partition of all samples is calculated as the average of the entropies of all clusters in the partition, weighted by the relative membership of each cluster. So the conditional entropy of cancer would be equal to the entropy of the resulting partition, i.e., *H*(*C*|*G*_1_,…,*G_n_*) = ∑*Ph*(*Q*). Then they calculated synergy as *H*(*C*) - *H*(*C*|*G*_1_,…,*G_n_*), where *H*(*C*) = *h* (number of cancerous samples/ total number of samples in the data).

#### Using mutual information to define candidate markers

In this work, we use single gene markers as a test bench to assess the value of distortion discovery and correction. To this end, we use mutual information to identify individual markers. We estimate mutual information from the genes’ continuous expression levels in the presence and absence of the phenotype of interest using an adaptive clustering algorithm as follows:

1. For each gene ***G_i_***
2. Sort the samples according to their value of ***G_i_***
3. Calculate mutual information **(I)** at each candidate decision boundary and save the best //candidate decision boundary = midpoint between each consecutive gene // values. #candidate decision boundaries = (n-1)
4. The algorithm terminates with the best two clusters for each gene that corresponds to best **(I)**
5. Candidate marker genes are the ones that have misclassified samples ɛ [MinMis,MaxMis]

MinMis and MaxMis stand for minimum/maximum number of misclassified samples of the candidate marker, respectively. They are two thresholds that are defined initially as 5% and 25% of the number of samples in the dataset, but could be slightly adjusted empirically. The purpose of using these two thresholds is to focus our search for candidate distorted markers, on the ones that have a considerable number of misclassifications; that might have some biological explanation. However, if the number of misclassifications are too small (for e.g. less than 5% of the number of samples) this could be counted as unexplainable noise. On the other hand, we do not expect the candidate distorted marker to have more than MaxMis misclassifications (eg. 25% of the samples) such that correction might not be useful or effective.

### 4.2.3 Defining spotters

A **spotter gene** is a gene whose behavior (e.g. expression level) is, ideally, not associated with the studied phenotype; so if distorted by hidden factors, it’s possible to define the affected samples if the spotter is significantly differentiated in them. Its value is when a marker is also sensitive to the same hidden factors causing mislabeling of the affected samples in its context. The spotter gene can be used to discover and correct for the hidden factors; hence reclaiming the marker’s prediction power.

### 4.2.4 Identifying candidate spotters

To trace down the effect of the hidden factors that is distorting a marker and causing mislabeling of some of its samples, we seek a spotter gene that is affected by the same hidden factors in the same set of affected samples. Being affected in those samples should mean that its value is differentiated (abnormally high or low) in those samples. We find the intersection of the marker’s mislabeled samples and the spotter’s outlier samples (in the tails of the probability density curve). We quantify the extent that the two genes (marker and spotter) have been affected by the same hidden factors using the Spotter’s Precision measure. This measure becomes maximum (1.0) when the spotter gene is not as differentiated in any other sample as it is in the affected samples. This happens when the tip of the tail is solely occupied by the distorted samples. Any sample that is not in the intersection but has a value, for the spotter gene, within the range of the affected samples should decrease the Spotter’s Precision measure. **Figure** shows the marker’s mislabeled samples that are spotted by the spotter; as they lie in the spotter’s tail and it shows the spotter’s spotting precision.

Mathematical model

We define the Spotter’s Precision with respect to a marker in a specific dataset (**Figure**) as follows:

Let 𝕊 denote the set of samples,

M ∈ [MinsMis, MaxMis] be the set of samples misclassified by the marker factor *F*^(*m*)^, *X*(*^F^*) be the value of the spotter factor *F*^(*s*)^ in sample 𝕤, *V*(*^D^*) ∈ [*high*, *low*] be the category of the spotter factor *F*^(*s*)^ in the set of distorted samples 𝕊^(*D*)^,

We define 𝕊^(*D*)^ as the set of distorted samples that represents the intersection of the marker’s misclassified samples and the samples in the spotter’s high or low tail (the tails are defined tentatively as Low-Tail ≤ μ-σ and High-Tail≥ μ+σ). The size of 𝕊^(*D*)^≥ MinMis and is defined as follows:

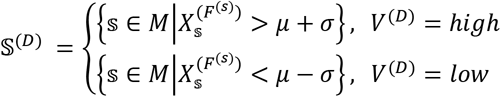

We define *τ* to be the set of samples in the spotter’s high or low tails (category = *high*/low) such that:

- For samples in the high tail, the spotter’s value is greater than or equal to its minimum value in the distorted samples 𝕊^(*D*)^.
- For samples in the low tail, the spotter’s value is less than or equal to its maximum value in the distorted samples 𝕊^(*D*)^.

The above condition favors the coherence of the affected samples in the tip of the spotter’s high/low tails. Hence, we define *τ* as follows:

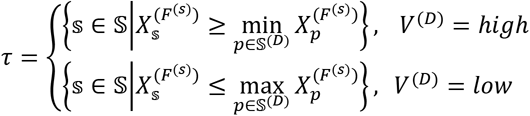

The Spotter’s Precision is a measure of how precisely the spotter spots the distorted samples. It is defined as the fraction of distorted samples to the samples in the spotter’s tail as follows:

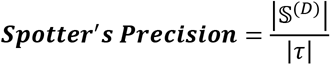

The Spotter’s Precision ∈ [0,1] is used to measure how precisely the spotter spots the distorted samples in the marker’s context. Such measure is used to score and rank distortion instances within a single dataset, and is later used as a threshold for selecting spotters for correction. **Table** lists the description of the used symbols.

### 4.2.5 Defining a distortion Instance

The distorted samples are the intersection of the mislabeled samples of the candidate distorted marker and the samples at one of the candidate spotter’s tails (category v ∈ V=high/low) such that they all have the same class (c ∈C=0/1). If the ratio of this intersection to all samples is significant; in the range [MinMis, MaxMis] then we count a new distortion instance, consisting of the distorted marker, and its misclassified samples with their correct class, the spotter and its category (**Figure**).

As a reminder, the use of the range [MinMis, MaxMis] is because we are looking for significant distortion instances; not too small to be counted as unexplained noise and not too large to be counted as a distorted marker, and such that correction would not help.

### 4.2.6 Modeling the relationship between a marker and its spotters

In this section we introduce a general model to analyze the relationship between a marker and its spotters. To model a distortion instance *D* representing a spotter factor *F*^(*s*)^ showing a category of abnormal values *v*∈*V*, used to capture the effect of a hidden factor on a marker factor *F*^(*m*)^ in a subset of distorted samples 𝕊^(*D*)^ of class *c*∈*C*; we use:

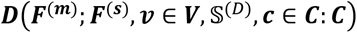

where ***C*** = {***c***_1_, ***c***_2_} is the set of actual classes of the distorted samples 𝕊^(*D*)^ and takes the values of ***c***_1_ = 0 or ***c***_2_ = 1.

***V*** = {*v*_1_, *v*_2_} is the set of category values of the spotter in the distorted samples 𝕊^(*D*)^ and takes the values of ***v***_1_ = *Low* or ***v***_2_ = *High*.

The above model describes a single distortion instance and it means that *F*^(*m*)^ is a marker (contains one feature or a set of features) that separates samples into clusters labeled ***c***_1_ and ***c***_2_ with a subset of distorted samples 𝕊^(*D*)^ of actual class ***c***. The predictive ability of *F*^(*m*)^ as a marker of *C* gets compromised as a result of distortion. An abnormal (*Low*/*High*) value *v* of the feature *F*^(*s*)^ in the similarly labeled *c* subset of samples(𝕊^(*D*)^), spots the cause of their mislabeling.

The above binary model can be generalized for ***C*** = {***c***_1_, ***c***_2_..***c_k_***} and ***V*** = {***v***_1_, ***v***_2_,.. ***v_n_***}, in order to model cases of more than two classes (e.g. cancer subtypes) and more feature categorical values, however the generalization is not straightforward^31^.

To model the cases when the marker is spotted in more than one distortion instance, we introduce a systemic relationship where a marker is spotted by a group of spotters, each, with its value ***v***, can explain a subset of the set of distorted samples 𝕊^(*D*)^ with actual class *c* in the context of that marker as follows:

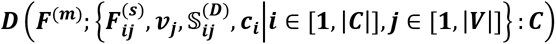

The above model means that *F*(*^m^*) is a marker; and most (or all) of its distorted samples, can be explained by a set of spotters 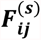, each with its noted category ***v***_j_, that are assumed to explain subsets of distorted samples each has one label ***c_i_***, where *i* ∈ [1, |*C*|] and *j* ∈ [1, |*V*|]. As a reminder,|*C*| = 2 and |*V*| = 2 in this study.

A more systemic relationship where a marker is spotted by a logical expression of spotting features in the form of a set of ANDed blocks that union up to cover most (or all) distorted samples in the context of the marker. Each block contains a set of ORed spotting features that spot the same mislabeling is shown below:

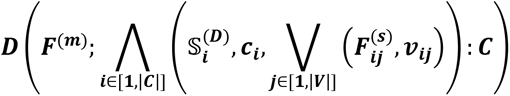

The above model means that ***F***^(m)^ is a marker whose distorted samples, can be explained by a set of ANDed tuples each contains a subset of distorted samples 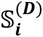 of class *c_i_* and a set of ORed tuples each contains a spotter 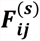 and its category *v_ij_*.

The previous model groups the distortion instances by class and category while the last model groups the distortion instances by the subset of distorted samples as well. This is useful for in-depth analysis as it aggregates all spotters spotting the same subset of the marker’s distorted sample and shows which subset of the distorted samples is spotted by which spotter(s).

#### Visualizing the relationship between a marker and its spotters

In **Figure 3** we show a hypothetical example of a synergistic gene pair marker (*fx,fy*) whose distorted samples are spotted by 7 spotter genes (*fi,fj*,*fk, fl*,*fm,fn*,*fo*). Solid lines represent Class (Red=**case**, Blue=**control**). Dashed lines represent Category (Red=**high**, Blue=**low**).

**Figure 3.**
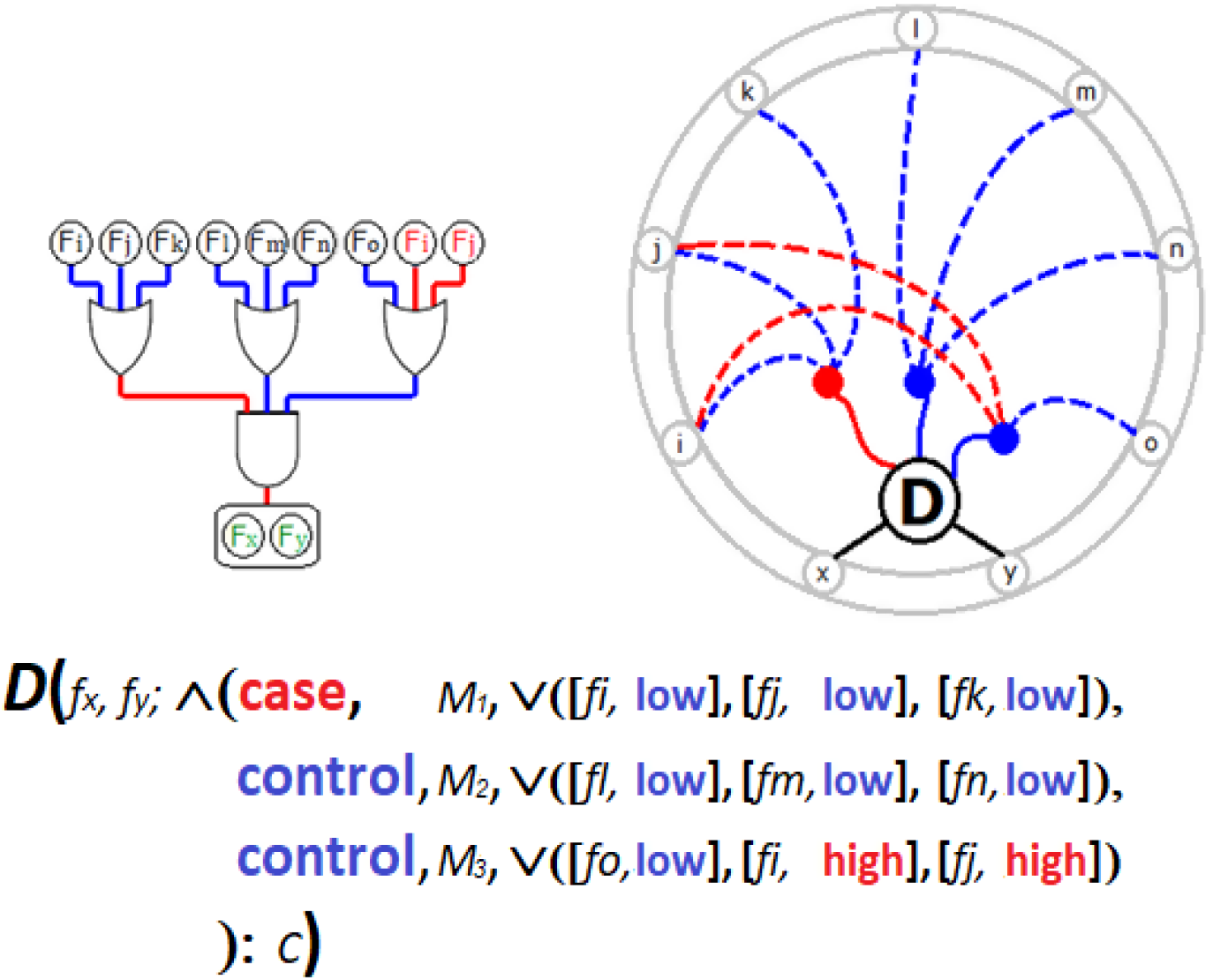
Spotter-Marker Relationship. A hypothetical example of a synergistic gene pair marker (*f_x_,f_y_*) whose distorted samples are spotted by 7 spotter genes (*f_i_,f_j_*,*f_k_, f_l_*,*f_m_,f_n_*,*f_o_*). Solid lines represent Class (Red=**case**, Blue=**control**). Dashed lines represent Category (Red=**high**, Blue=**low**).

**Figure** shows an actual example of a synergistic gene pair marker (RBP1, HLA_DPB1) whose distorted samples are spotted by nine spotter genes (ACSM1, SIM2, TPSAB1, OLFM4, AGR2, ERG, COMP, CFD, CALD1). For example, the false positive mislabeled samples (**N46,N43, N38,N19,N23,N18**) are explained either by the down-regulated gene ***TPSAB1*** or the down-regulated gene ***OLFM4*** or the down-regulated gene ***AGR2*** or the up-regulated gene ***ERG***.

### 4.2.7 Distortion discovery and correction

The following algorithm searches for distortion instances in the datasets. A distortion instance is characterized by a spotter gene abnormally low or highly expressed in a group of samples that are mislabeled in the marker’s context, while the behavior of the spotter gene is undifferentiated in the rest of the samples.

***Input***: The algorithm takes an omics dataset in the form of a matrix of rows by columns. Each row represents individual gene values across samples, and each column represents the genetic profile of a sample and the sample’s label.

***Output***: The algorithm outputs distortion instances associated with a distortion quality *score (score = spotter’s precision)*. Each distortion instance consists of a *spotter* gene that is low or highly expressed (c*ategory*), its corresponding set of false positive or false negative *distorted samples* and the correct *class* in the context of a *marker* gene, the distorted marker’s initial information with the class, the marker’s corrected information and the synergy between the marker and spotter with respect to the class. The mutual information and pairwise synergy tables of all the genes in the dataset are also included in the output. This is the design of the distortion record, in case of single gene marker (1) and multiple gene markers (2):

1. (Fs, Fm, Class, Category, Score, I, I_Cor, Syn-2, {Distorted Samples}).
2. (Fs, {Fm}, Class, Category, Score, Syn, Syn_Cor, Syn-n, {Distorted Samples}).

#### Distortion Discovery Algorithm

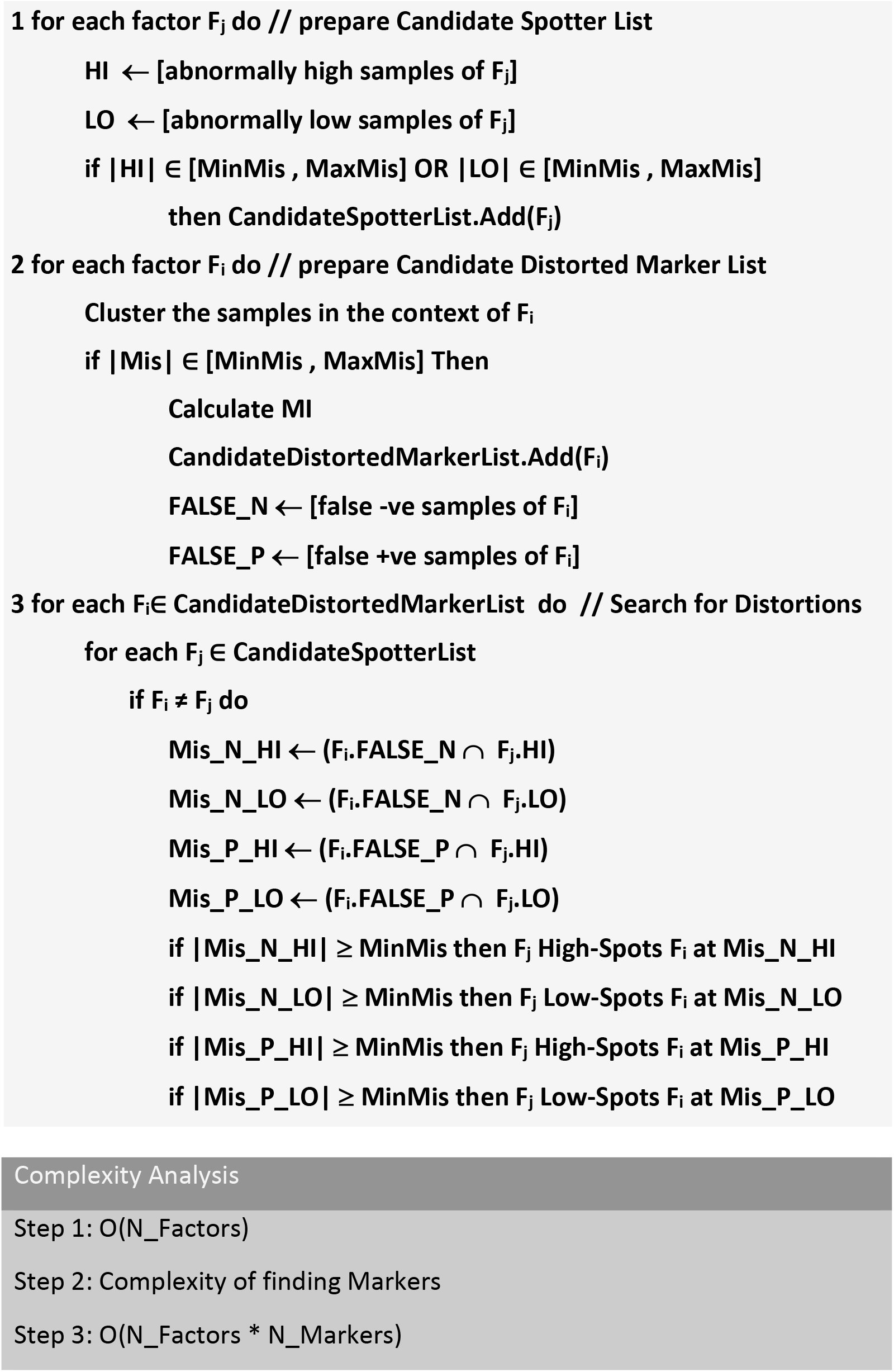

The complexity of the algorithm depends on the marker cardinality; # single markers = N_Factors while # marker pairs = N_Factors^2^/2. In this application, N_Factors is number of features in the input dataset.

#### Correcting distortion effect on a marker

The objective of this section is to explain how correction for the effect of distortion is done during prediction. First, we explain the output of the training process, then we define some terms used in correction and testing, and finally we describe the correction algorithm.

Distortion instances are discovered during the training process, using the Distortion Discovery algorithm described above. We train our model to learn the impact of distortion on the relationship between the distorted marker and the spotters. After training, we get a trained machine that has learned which markers are distorted and which are not, according to our previously defined criteria, as well as the relationship between each distorted marker and its spotter(s).

The output of the training process is a list of predictors (***PLIST***). ***PLIST*** is a list of predictor records defined as: **P struct** {Marker, Class, Category, Decision Boundary, Prediction Rate, Number of Spotters, List of ***Spotter***} where each ***Spotter*** is a **struct** {Spotter Gene, Spotter Tails}. The list of predictors is then deposited in the Distortion Repository, to be used for prediction testing.

This means that the input to the prediction testing will be a list of predictors, each predictor is a set that consists of the distorted marker, the class of its distorted samples and the categorical value of its spotters, the marker’s decision boundary, its prediction rate as defined below, number of spotters and a list of the spotters. The spotter consists of a gene and its tails – used for the purpose of cross dataset predictions.

#### Definitions

*MarkerPredictionRate* : the ratio of the number of times the marker was involved in prediction decisions to the total number of training iterations, used in the correction algorithm as the marker’s voting weight.

*SpottingRate* : The ratio of polling spotters to the total number of spotters for every predictor.

MinSpottingRate : A threshold for distortion spotting credibility.

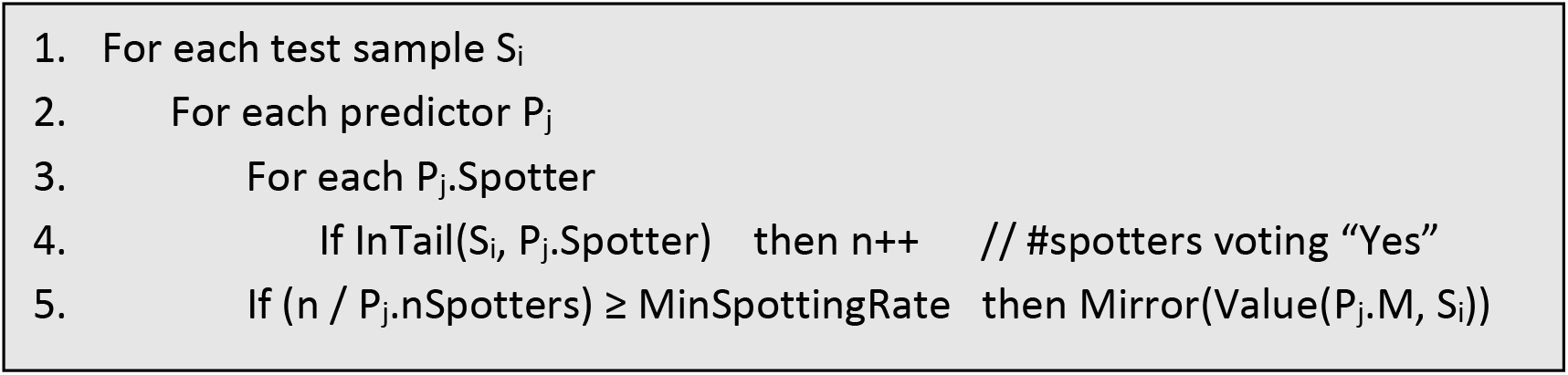

#### Correction Algorithm

The above correction is done for each sample during prediction testing. Note that the correction is done by flipping the class label of the sample, or more generally mirroring the sample’s classified position across the decision boundary. This is only done, if there is enough number of spotters voting for the correction. That’s to say, if the number of polling spotters are greater than or equal to a predefined

*MinSpottingRate* threshold.

### 4.2.8 Assessment

#### Assessment of Significance

Permutation tests for datasets

For statistical validation, we implement permutations on each dataset, whose rows correspond to genes and columns correspond to samples. In each permutation, the columns are randomly shuffled in order to dissociate the gene profile of each sample with its class label (e.g. case vs control), while maintaining the genetic profile of each sample. We permute the dataset *nPerm* times and discover distortions. Then we count the number of permuted datasets, with number of distortions scoring greater than or equal *TopScore,* that were greater than or equal to that of the actual dataset. Then we divide that number by *nPerm* to calculate p-value. A small p-value means that the high scored distortion instances have a small probability of happening by chance.

We compare the frequancy distribution of the distortion instances scores of the actual dataset to the frequency distribution of the average distortion instances scores of all permuted datasets (1000 permutations). *TopScore* is then selected as the lowest possible score with lowest p-value.

#### Permutation tests for assessing the significance of cumulative overlap function

We create a pool of genes from each dataset, then randomly create distortion instances equal in size to that of the actual distortion instances in each dataset. Then a thousand permutations (with replacement) are done, with a count of the number of permutations that produce cumulative overlap function value similar to the actual. The null hypothesis tested by this permutation test is that the cumulative overlap for the actual distortion instances can be produced by chance using similar number of randomly permuted distortion instances for each dataset.

#### Permutation tests for assessing the significance of overlap with immune/stromal signatures

For each dataset, we first remove all genes from the signature that are not in the dataset (***Modified_Signature***), and we randomly draw a set of genes ***G*** from the dataset equal in size to that of the actual spotters identified from it and compute the overlap with the ***Modified_Signature*** (we do this once for immune and another for Stromal signatures). We repeat a thousand permutations and calculate p-value at the end. (p-value = number of times null hypothesis was true/1000). The null hypothesis tested by this permutation test is that the number of permuted spotters’ overlap with the signature is greater than or equal to the number of actual spotters’ overlap with the signature. (Note that the probability of random overlap per dataset = |***G*| / |Dataset| x |*Modified_Signature*|**). This is also repeated for the total top spotters identified from all eight datasets, as well as the network of top spotters identified from all eight datasets, after creating a pool of genes from the eight datasets. In the case of top spotters network permutations testing, the randomly selected gene sets, that are each equal in size to the actual set identified in the network of top spotters, are repeatedly (1000 times) fed into a network analysis algorithm, and the resulting networks are assessed for overlap.

#### Assessment of utility

##### Prediction tests

Prediction testing is done to assess the ability of the Spotter-Marker model in improving the predictive performance of classifiers. For this purpose, we utilize two settings:

1. Cross validation: For each dataset, 90% of the samples are randomly selected for training, and the remaining 10% are used for testing. This is done 1000 times (with replacement with respect to the 1000 trials).
2. Cross dataset validation: Performed by testing the trained model on independent datasets.

The markers used for testing are the ones found to have top corrected mutual information (**MI**) with the phenotype, during training, and that pass a minimum threshold i.e. their MI is greater than the minimum corrected mutual information (**MinICorr**). The markers that possess such criteria are used together with their spotters for prediction testing. The test sample is checked at each of the tails of all the marker’s spotters. If it lies in the tail of a minimum number of them defined by a threshold (*MinSpottingRate*), then we flip the predicted class of the sample, or generally speaking, we mirror the position of the point across the decision boundary of the marker i.e. move it to the cluster where it belongs on the opposite side of the decision boundary as follows:

**Mirror(DecisionBoundary, X) { return DecisionBoundary+DecisionBoundary-X; }**

We define several decision boundaries across each marker’s value range, representing a moving threshold to compute the Receiver Operating Characteristic (ROC) curve. Each test sample is classified using each decision boundary (predicted class). Each predicted class is compared with the actual class of the test sample and one of the four counters (True Positive (*TP*). True Negative ^49^, False Positive (*FP*). False Negative ^50^) is incremented -by adding *MarkerPredictionRate* - accordingly. Incrementing by *MarkerPredictionRate* involves the weight of the marker in the voting.

This is done once without the decision of the markers’ spotters (without correction) and once including the decision of the spotters (with correction). This is also repeated a third time for the top non-distorted markers (their MI > minI) in the dataset. Three ROC curves are juxtaposed comparing the results of the three prediction methods.

Sensitivity and specificity are calculated from true and false positive and negative rates. Where true positive rate (TPR) is defined as the ratio of the number of correct positive predictions to the total number of positives, and false positive rate (FPR) is defined as ratio of the number of incorrect positive predictions to the total number of negatives, Sensitivity = TPR, Specificity = 1 – FPR. Then the areas under the ROC curves (AUC) are computed, for the curve that corresponds to prediction results without correction, for the curve that corresponds to prediction results after using correction and for the curve that corresponds to prediction results done with non-distorted markers. The latter are the top markers in the dataset defined without involving the concept of distortion.

#### Assessment of reproducibility

##### Prediction Testing on validation datasets

In order to assess the reproducibility of our prediction results, we test prediction on validation datasets other than the training datasets, the list of predictors ***PList*** (

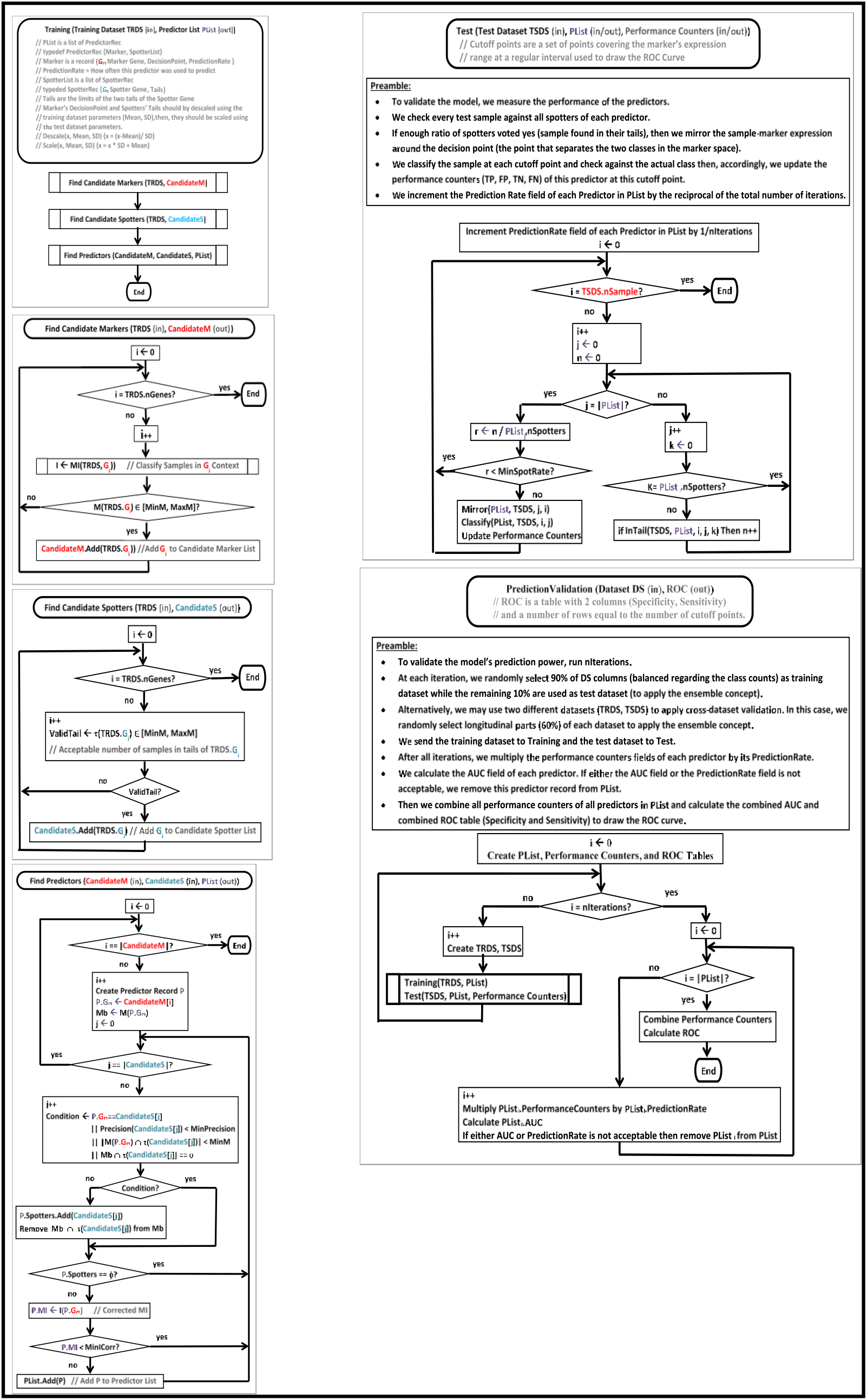

) which is created during training, is stored in the Distortion Repository (in a dataset independent format). Gene standardization, with respect to normal distribution, a location-scale method for batch effect removal ^3^, is used to translate the gene values across datasets. Standardizing the Marker’s DecisionBoundary is done by subtracting the mean of the marker’s expression level and dividing by its standard deviation before saving it in the Distortion Repository (*descaling*). Each spotter’s percentile (the start of the tail) is also standardized using the mean and standard deviation of the expression of the spotter gene. Then before using them on a validation dataset, we do the reverse, i.e. multiply the marker’s decision boundary by the standard deviation of the validation dataset, and add to it the mean of the validation dataset (*scaling*). The same is done with the spotters’ tail(s).

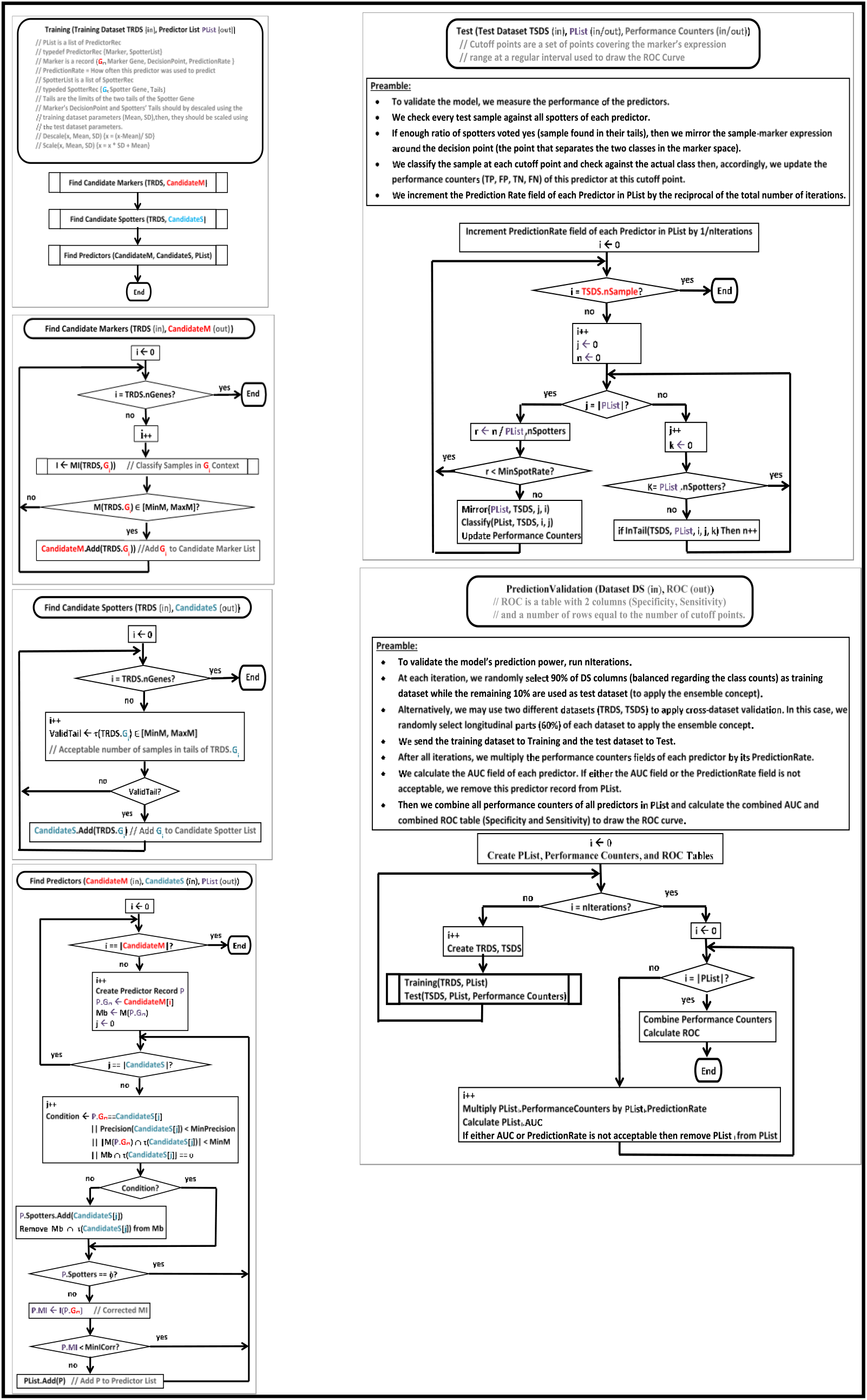

shows the modules and flowchart of Prediction Validation.

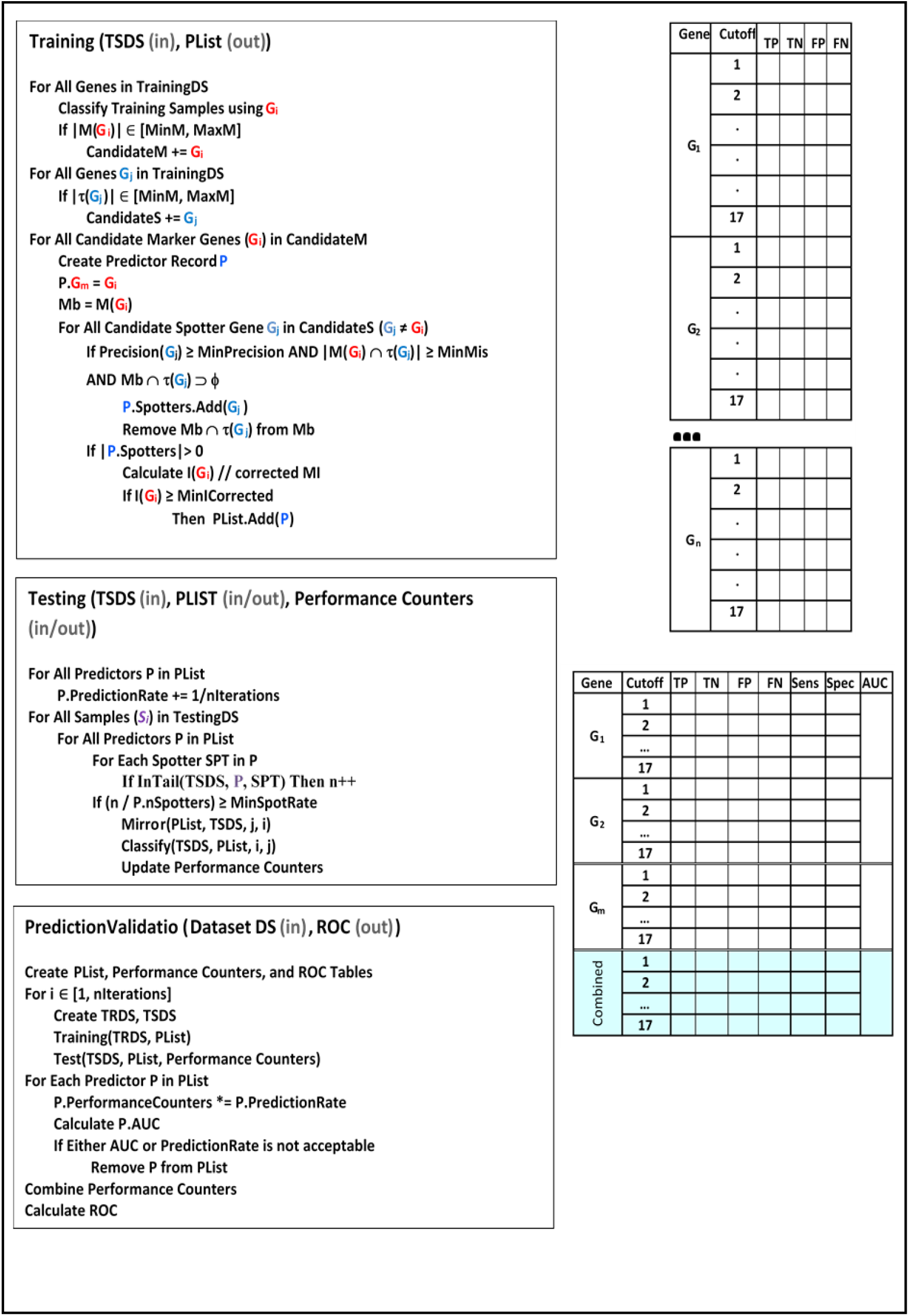

#### Assessment of Biological Relevance

##### Overlap analysis

Overlap analysis is done in order to assess the biological relevance of the distortion instances discovered on different datasets. It is done on the statistically significant top scoring distortion instances across the studied datasets.

Overlap Matrix:

Each overlap matrix is a *k*×*k* matrix that represents the pairwise overlap between the significant spotters, markers or spotter-marker pairs in each pair of datasets. We define the Overlap Ratio ***Γ ij*** between two datasets *i* and *j*, as the Jaccard coefficient of the overlapping genes (spotters/markers /spotter-marker pairs) between two datasets. That’s to say, the fraction of common significant genes (spotters/markers/spotter-marker pairs) in the two datasets among the number of significant genes (spotters/markers/spotter-marker pairs) in the two datasets. The Overlap Ratio is defined as:

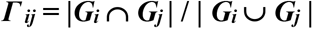

where *G_i_* and *G*_j_ denote the sets of significant genes such that G=F_s_ in case of assessing the overlap between spotter genes across datasets *i* and *j*, respectively, G= Fm in case of assessing the overlap between marker genes across datasets *i* and *j*, respectively and G = (F_s_, F_m_) pair in case of assessing the overlap between spotter-marker instances across datasets *i* and *j*, respectively.

**Cumulative Overlap Function:**

A cumulative overlap function is defined in order to quantify the overall overlap between the *k* datasets representing identical phenotype classes, e.g. datasets that have the two classes “prostate tumor” and “normal prostate” are compared together. The cumulative overlap function is a function in the form f:{1,…, k} → [0,1], assessing the fraction of genes, either markers, spotters, or spotter-marker pairs that are discovered to be significant in at least a given number of datasets of the same biology. The *cumulative overlap* function *γ(l)* for 1 ≤ *l* ≤ *k* is defined as:

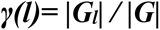

where *Gl* denotes the set of genes (spotters/markers/spotter-marker pairs) that are found to be significant in at least *l* datasets of the same biology. And **G** is the union of all significant genes in all datasets of the same biology. Observe that 0 ≤ *f*(*i)* ≤ *1, f*(*1) =1*, and *f*(*i)* is a monotonically non-increasing function of *i*.

##### In-Depth Gene Set Analysis

This is a per dataset analysis where all distortion instances of each dataset are searched for overlap on common distorted samples, allowing partial overlap. For each overlapping group of distortion instances, we determine the union of their spotter genes, the union of their marker genes and the union of the two unions. We select the overlapping groups with the largest number of common distorted samples. The different unions of genes are used in gene set enrichment analysis (GSEA) and network analysis. The purpose of this is to assess common biological mechanisms, if any, underlying the distortion of those common distorted samples.

### 4.2.9 Development and Execution Environment

The software is developed in visual C++ v12-x64 on Windows 10 using standard libraries. It runs in parallel and uses n-1 of the available cores on the pc. It takes 1 to 3 hours to run 1000 permutation iterations on an 8-core 4G Hz Core i7, hence the permutations and predictions modules were run in several sessions on several pcs.

## 4.3 RESULTS

In this section, we present the validation results using eight gene expression datasets. We first validate the statistical significance of the results, then we demonstrate the gain in distorted markers’ mutual information in all datasets, we also demonstrate how some strong marker genes show weak mutual information with the phenotype of interest in one dataset, but regain its predictive ability after discovery and correction. The same is also demonstrated for the markers of the phenotype of interest that are already established in literature, that would show low mutual information with the disease in some datasets of the same biology of interest, due to the effect of distortion, but reclaim their high mutual information with the disease after discovery and correction. We also show prediction cross-validation results on the datasets as well as prediction performance on four validation datasets to assess the utility and reproducibility of the introduced model. Then we compare the performance of the Spotter-Marker model to that of other machine learning algorithms, in order to assess the value of the Spotter-Marker model. Finally, we discuss the biological relevance of the spotters, markers and distortion instances, supported by the overlap analysis of the discovered spotters, markers and the distortion instances across datasets of the same phenotype of interest.

### 4.3.1 Datasets of Prostate Cancer

We use eight publically available prostate cancer datasets (case/control, subtypes, metastatic/non-metastatic, aggressive/non-aggressive) representing different types of omics data. The datasets are described in **Table 2**. DS1 ^32^ and DS2 ^33^ are microarray expression profiling for prostate tumor and control samples. DS3 ^34^ is an RNA sequencing expression for samples belonging to one of two most common prostate cancer molecular subtypes (ERG, ETV1). DS4 ^35^ and DS7 ^33^ are microarray expression profiling for metastatic prostate tumor and non-metastatic prostate samples. DS5 ^36^ is an RNA sequencing expression for treatment-emergent small-cell neuroendocrine prostate cancer (t-SCNC) and Adenocarcinoma samples. DS6 ^33^ is microarray expression profiling for samples with metastatic prostate tumor and control samples. DS8 ^37^ represent genome-wide DNA methylation data for prostate tumor and benign prostate samples.

**Table 1.** Description of the symbols used in the Spotter-Marker mathematical model.

| Symbol | Description |
| --- | --- |
| $\mathcal{S}$ | The set of all samples in the dataset |
| $X$ | The set of values ($X_s^i$ = value of gene $i$ in sample $s$) |
| $V$ | Category of abnormality (high / low) |
| $D$ | Distortion ($\mathcal{S}^D$ = Distorted Samples ) |
| $F$ | Factor ($F^m$=marker factor, $F^s$=spotter factor) |
| $C$ | Class (case / control) |
| $M$ | Set of mislabeled samples |
| $\tau$ | The set of samples in the tail (where the spotter gene has abnormal value) |
| $\mu, \sigma$ | Mean and Standard Deviation of the spotter's values in all samples |

**Table 2.** Description of the datasets used in this study. The dataset name, type, platform, number of samples, number of features, class labels and date of creation.

| Dataset | Type | Platform | # Samples | # Features | Class Labels | Date |
| --- | --- | --- | --- | --- | --- | --- |
| DS1 | Prostate Cancer microarray Expression profiling | Affymetrix HG-U95Av2 | 102 Samples: 52 prostate tumor, 50 non-tumor | 12,625 | Prostate Tumor/Control | 2002 |
| DS2 | Prostate Cancer microarray Expression profiling | Agilent-014698 HG CGH Microarray 105A (G4412A) | 56 Samples; 28 prostate tumor, 28 non-tumor | 11,131 | Prostate Tumor/Control | 2012 |
| DS3 | Prostate Cancer RNA-seq Expression | IlluminaGA and IlluminaHiseq RNASeq | 50 Samples: 25 Subtype ERG, 25 Subtype ETV1 | 17,427 | Prostate Cancer Subtypes ERG/ETV1 | 2015 |
| DS4 | Prostate Cancer microarray Expression profiling | Almac Diagnostics Prostate Disease Specific Array (DSA) | 112 Samples: 56 Non-Metastatic, 56 Metastatic | 15,096 | Metastatic Prostate Tumor/ Non-Metastatic Prostate | 2018 |
| DS5 | Prostate Cancer RNA-seq Expression | IlluminaGA and IlluminaHiseq RNASeq | 28 Samples: 14 Adenocarcinoma, 14 t-SCNC | 18,538 | t-SCNC/ Adenocarcinoma | 2018 |
| DS6 | Prostate Cancer microarray Expression profiling | Agilent-014698 HG CGH Microarray 105A (G4412A) | 56 Samples; 28 metastatic castrate resistant prostate tumor, 28 Normal Prostate | 11131 | Metastatic Tumor/Control | 2012 |
| DS7 | Prostate Cancer microarray Expression profiling | Agilent-014698 HG CGH Microarray 105A (G4412A) | 64 Samples: 32 Primary prostate tumor, 32 metastatic castrate resistant prostate tumor | 11131 | Metastatic Prostate Tumor/ Non-Metastatic Prostate | 2012 |
| DS8 | Prostate Cancer genome-wide DNA methylation | Illumina Infinium 450K Human DNA Methylation Beadchip | 126 Samples: 63 Benign Prostate, 63 Tumor Prostate | 8546 | Benign Prostate / Tumor Prostate | 2017 |

**Table 3.**
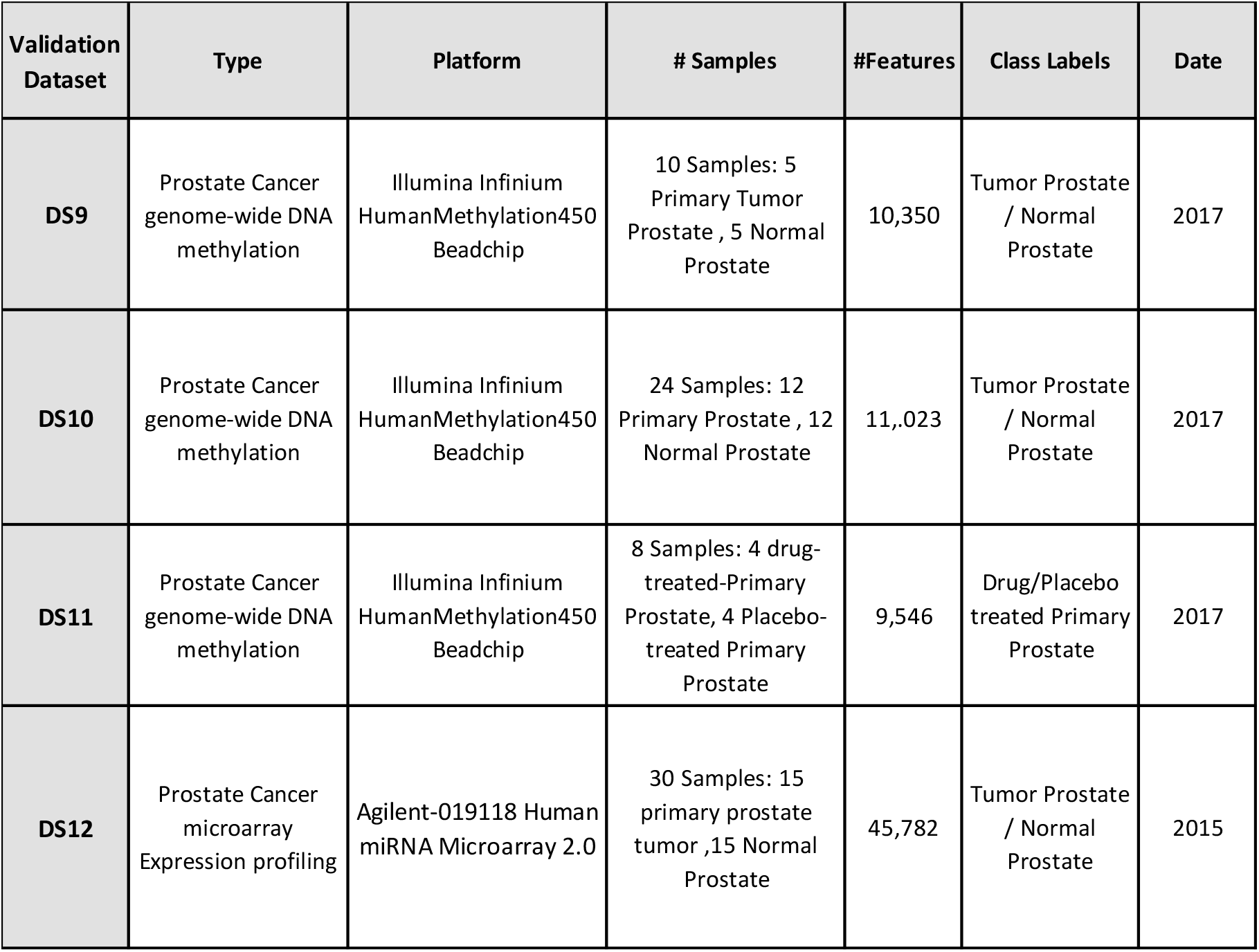
Description of the validation datasets used in this study. The dataset name, type, platform, number of samples, number of features, class labels and date of creation.

| Validation Dataset | Type | Platform | # Samples | #Features | Class Labels | Date |
| --- | --- | --- | --- | --- | --- | --- |
| **DS9** | Prostate Cancer genome-wide DNA methylation | Illumina Infinium HumanMethylation450 Beadchip | 10 Samples: 5 Primary Tumor Prostate , 5 Normal Prostate | 10,350 | Tumor Prostate / Normal Prostate | 2017 |
| **DS10** | Prostate Cancer genome-wide DNA methylation | Illumina Infinium HumanMethylation450 Beadchip | 24 Samples: 12 Primary Prostate , 12 Normal Prostate | 11,023 | Tumor Prostate / Normal Prostate | 2017 |
| **DS11** | Prostate Cancer genome-wide DNA methylation | Illumina Infinium HumanMethylation450 Beadchip | 8 Samples: 4 drug-treated-Primary Prostate, 4 Placebo-treated Primary Prostate | 9,546 | Drug/Placebo treated Primary Prostate | 2017 |
| **DS12** | Prostate Cancer microarray Expression profiling | Agilent-019118 Human miRNA Microarray 2.0 | 30 Samples: 15 primary prostate tumor ,15 Normal Prostate | 45,782 | Tumor Prostate / Normal Prostate | 2015 |

#### Validation Datasets

For prediction testing on validation datasets, we use four publicly available prostate cancer expression datasets. Two datasets DS9 ^38^, DS10 ^39^ are DNA methylation data for prostate tumor and benign prostate samples. The third dataset DS11 ^40^ is DNA methylation data for drug-treated primary prostate tumor and placebo-treated primary prostate tumor samples. The fourth dataset DS12^41^ is a microarray expression profiling for prostate tumor and control samples (**Table**).

### 4.3.2 Statistical significance of Results

The eight plots in Error! Reference source not found. show the results of permutation testing on the eight datasets. In each plot, the red and blue cumulative frequency curves represent the distortion scores found in the actual dataset versus the average of the distortion scores found in a thousand permuted datasets, respectively. The plots show that the actual datasets have significantly higher frequencies of the top scores as compared to the permuted ones. Top scores are defined by a threshold of statistical significance; a cut-off score above which we have the most confidence in rejecting the null hypothesis. They are the lowest possible scores of the distortion instances in each dataset, that are chosen to have the lowest p-value. The table shows the p-values of the selected top scores in each dataset. The top scores represent the scores of the significant top distortion instances selected from each dataset for further analysis.

The eight plots in Error! Reference source not found., each represents one of the eight datasets, demonstrate the distortion scores versus spotter-marker synergy. All eight plots are in concordance and show that the high distortion scores characterize the zero and very low synergy values between spotter and marker pairs. This indicates that the higher scoring spotter-marker pair of genes, tend to have very low to no direct – cooperative or redundant – relation with each other, and also suggests, that the spotter is not a marker, itself, of the phenotype of interest.

### 4.3.3 Gain in mutual information

**Figure** demonstrates, the quantitative gain in mutual information (MI) in the studied datasets. The figure shows the top improvements in MI in the eight datasets. For example, the top plot shows marker genes with corrected MI ≥ 0.8. Each vertical line starts from the minimum MI found for a gene, and ends at the maximum MI found for this gene in all datasets. Established Prostate Cancer markers are given a different color to be easily distinguished from other genes. For each gene, the bar starts at initial MI before correction and ends at the maximum corrected MI. The three plots show that almost all genes were corrected to more than their maximum MI found in all datasets.

### 4.3.4 Comparison of results on different datasets

Error! Reference source not found. demonstrates the following scenario: *Gene Gm is a good marker in DSx, is Bad in DSy, and was corrected back to good in DSy*. Which means that a gene which is a marker in one dataset *x* – or an established marker of the biology of interest- was not showing up as marker in another dataset *y* of the same biology, due to its low mutual information with the biology of interest in the dataset *y*. Such gene was reclaimed as marker of the same biology after discovery and correction in dataset *y*. For example, **MYC** is a known prostate cancer marker ^42^, that had a low MI with the disease in all datasets, but was corrected with a maximum correction of 0.665, hence reclaiming its predictive power. This scenario shows another validation of the model’s reproducibility.

### 4.3.5 Prediction Performance

In the following tests, the false positive predictions were less spot-able. Therefore, the software was set to focus on correcting for false negative predictions. Top markers are used to draw ROC curves. They are defined as the predictors that, when used, result in the highest AUC in the Prediction Test. Top markers are selected such that in each dataset, no other markers result in AUC equal to or greater than their minimum AUC. We try to choose a balanced number of distorted and non-distorted top markers for the sake of comparison of both results.

#### Cross Validation

The cross validation results are shown in **Figure 30**. The eight plots portray an improved predictive ability of the corrected markers in the eight datasets. This is represented by the improved AUC after correction as opposed to the AUC before correction in each dataset. The plots also show that the predictive ability of the corrected markers are comparable, or better, to the perfect, undistorted markers in all datasets.

The plot of the first dataset (DS1), for example, indicates a prediction improvement demonstrated by an increase in the AUC from 0.559 before correction to 0.865 using the top 243 corrected markers (Top markers are chosen with selection threshold AUC>=0.85). In DS1, there are eight undistorted markers (with AUC>=0.85) only. The corrected markers perform better than the non-distorted markers whose resulting AUC is 0.825. Another example is the fourth dataset (DS4), there is an improvement in the corrected markers’ predictive ability demonstrated by the increase in AUC from 0.556 before correction to 0.864 after correction using almost a balanced number of distorted and non-distorted markers (18,20). Moreover, the corrected markers perform much better than the non-distorted markers, the latter has AUC = 0.584 as shown **Figure 4**.

**Figure 4.**
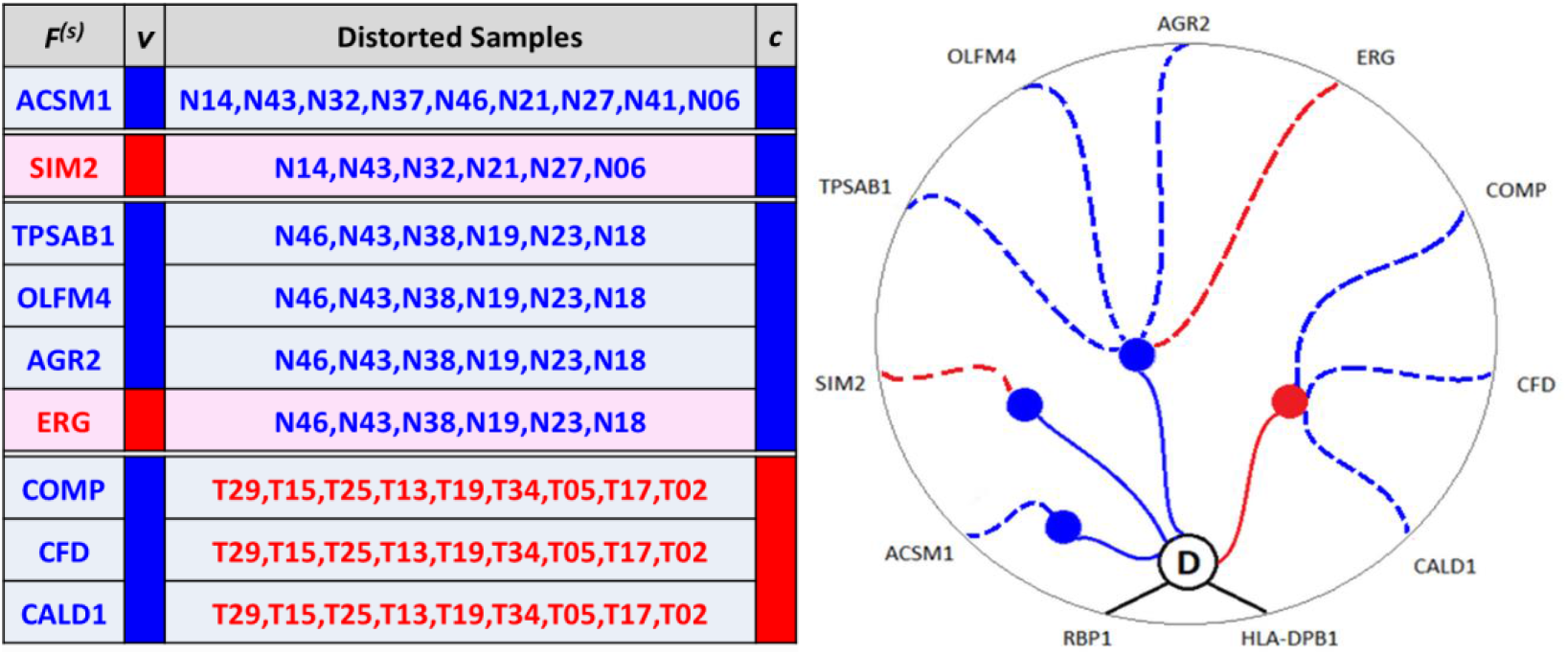
An Example of Spotter-Marker Pair Relationship. The synergistic gene pair marker (RBP1, HLA_DPB1) whose distorted samples are spotted by nine spotter genes (ACSM1, SIM2, TPSAB1, OLFM4, AGR2, ERG, COMP, CFD, CALD1). The false negative subset of the marker’s distorted samples (**T29,T15,T25,T13,T19,T34,T05,T17,T02**) are spotted by either the down-regulated gene ***COMP*** or the down-regulated gene ***CFD*** or the down-regulated gene ***CALD1*.** While the false positive subset of the marker’s distorted samples (**N46,N43,N38,N19,N23,N18**) are spotted either by the down-regulated gene ***TPSAB1*** or the down-regulated gene ***OLFM4*** or the down-regulated gene ***AGR2*** or the up-regulated gene ***ERG*.**

**Figure 5.**
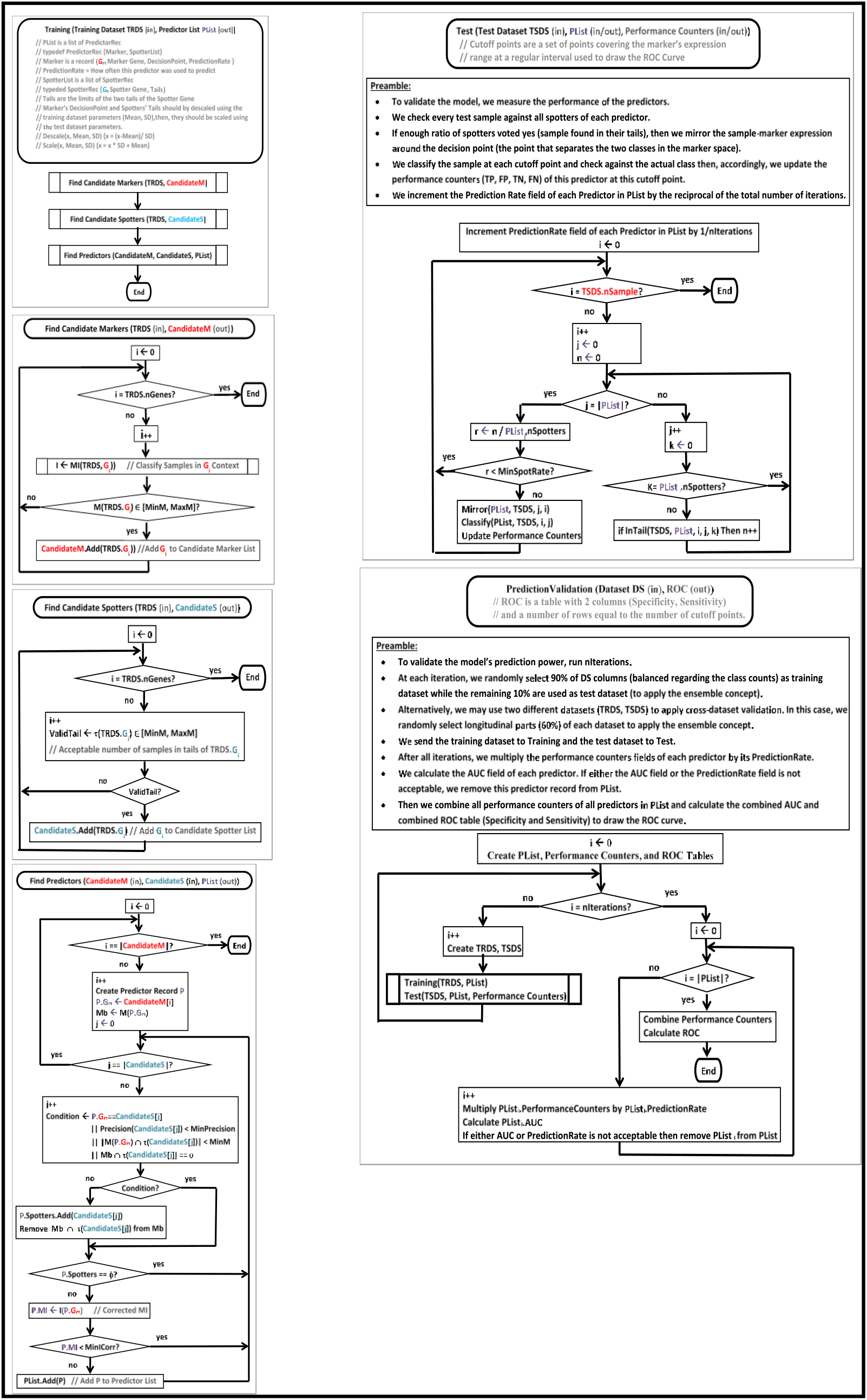
Prediction Validation. Pseudocode (upper panel) and flowchart (lower panel)

**Figure 6.**
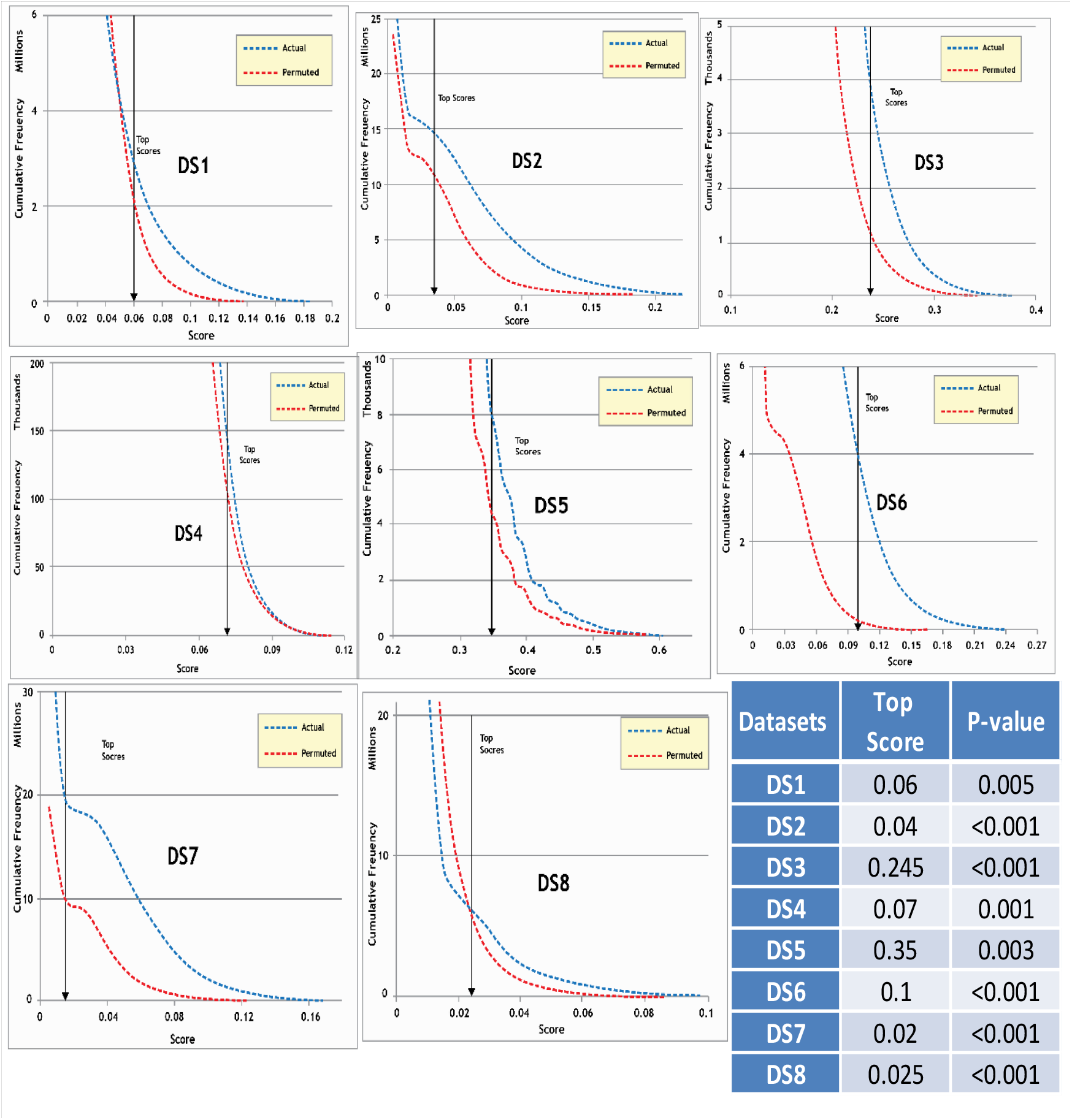
Statistical Significance of Results. The figure shows the results of permutation testing on the eight datasets. The blue curve in each plot represents the frequency distribution of the scores in the actual dataset and the red curve in each plot represents that of the average scores over a thousand permuted datasets. The table shows the p-values of the selected top scores in each dataset. The top scores represent the scores of the significant top distortion instances selected from each dataset for further analysis.

**Figure 7.**
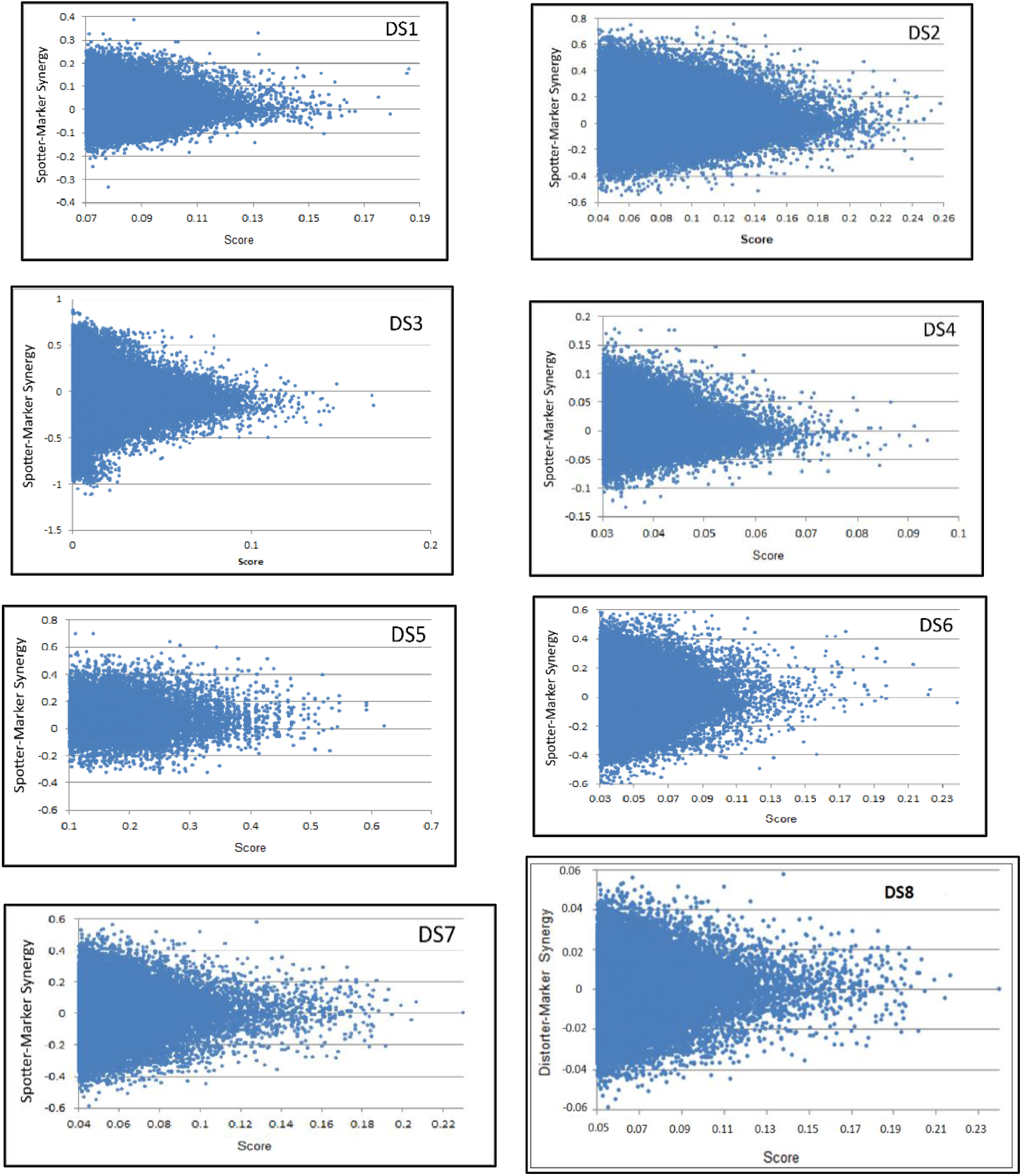
Spotter-Marker Synergy. The eight plots show the synergy between the Spotter-Marker gene pair versus the Spotting Score (Spotter’s Precision) in each of the 8 datasets. All plots show that score is maximum around zero synergy.

**Figure 8.**
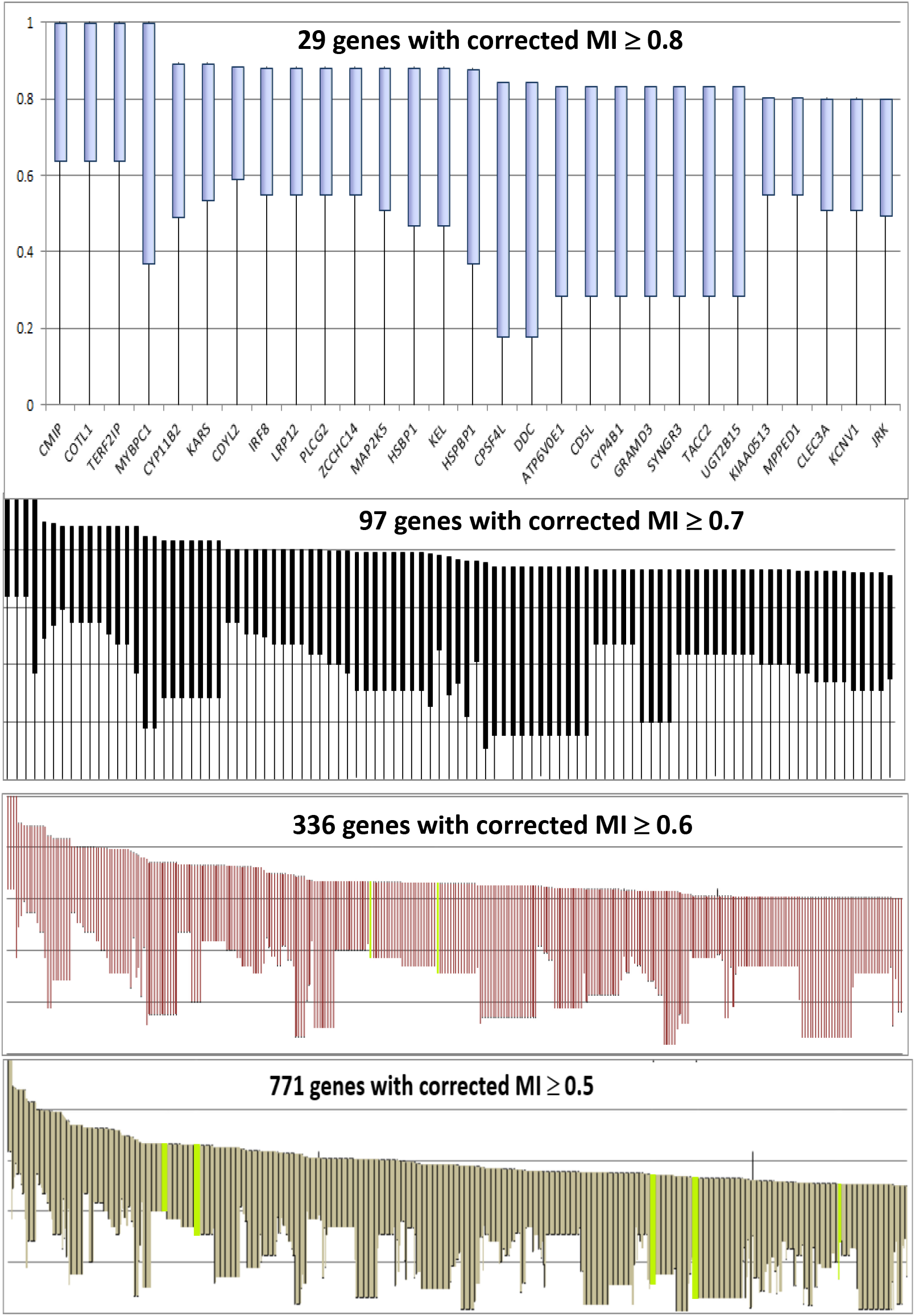
Gain in Mutual Information. This figure shows the top genes’ gain in mutual information (MI) with the biology of interest. For example, the top plot shows marker genes with corrected MI ≥ 0.8. The line starts from the minimum MI found for this gene in all datasets, and ends at the maximum MI found for this gene in all datasets. Established markers are given a different color to be easily distinguished. For each gene, the bar starts at initial MI before correction and ends at Corrected MI. All genes

**Figure 9.**
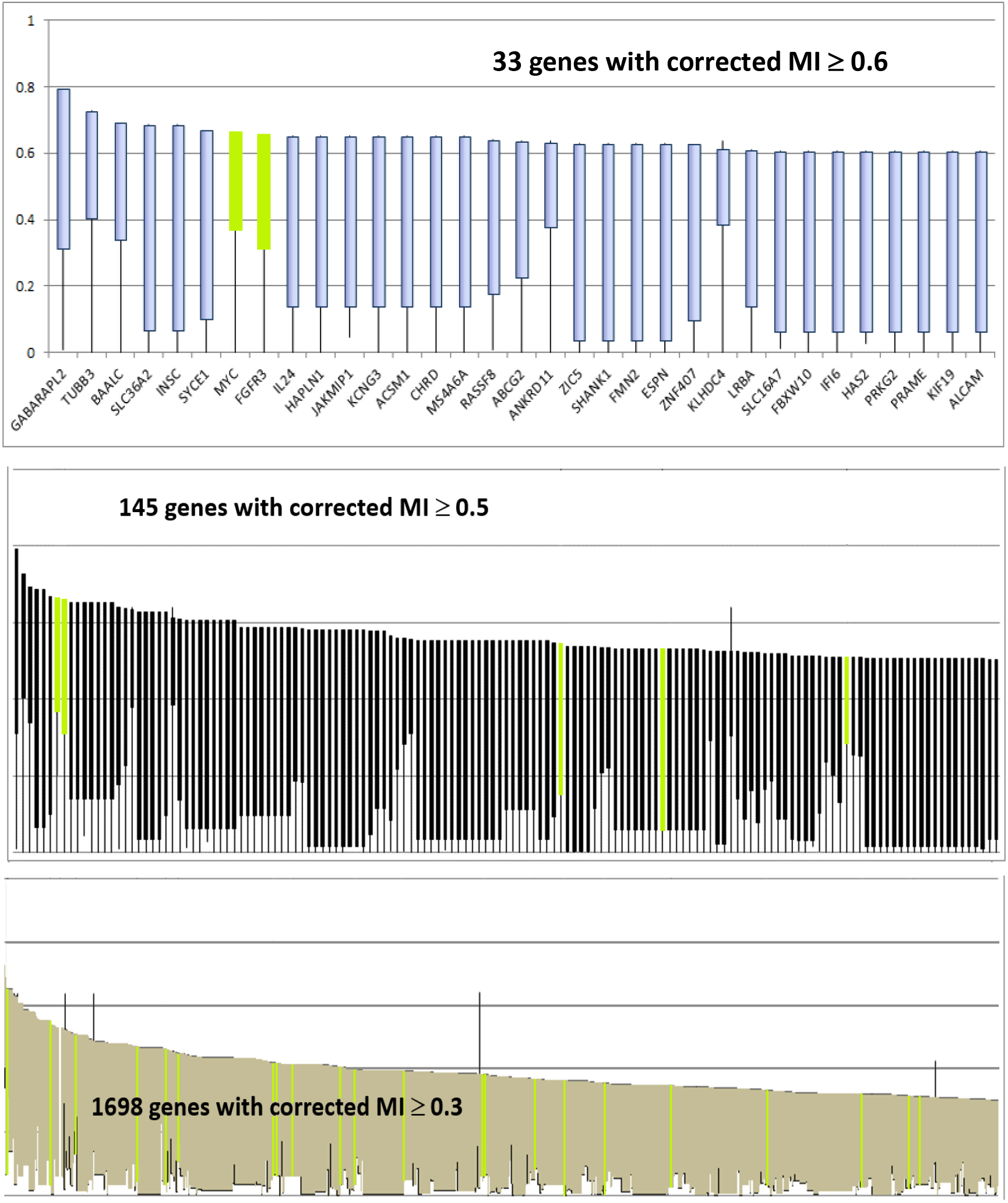
Comparison of results on different datasets. Marker genes with corrected Mutual Information(MI) ≥ 0.6, the line starts from the minimum MI found for this gene in all datasets, and ends at the maximum MI found for this gene (not where it was corrected). Established markers of the biology of interest are given a different color to be easily distinguished. For each gene, the bar starts at initial MI before correction and ends at Corrected MI. All the above genes, are either global markers, or had high MI in another dataset, *not where it was corrected.* All genes were corrected to more than (or very close to) their maximum MI found in all datasets. The diagram demonstrates the scenario: *Gene G_m_ is a good marker in DSx, is bad in DSy, and was corrected back to good in Dsy*.

**Figure 10.**
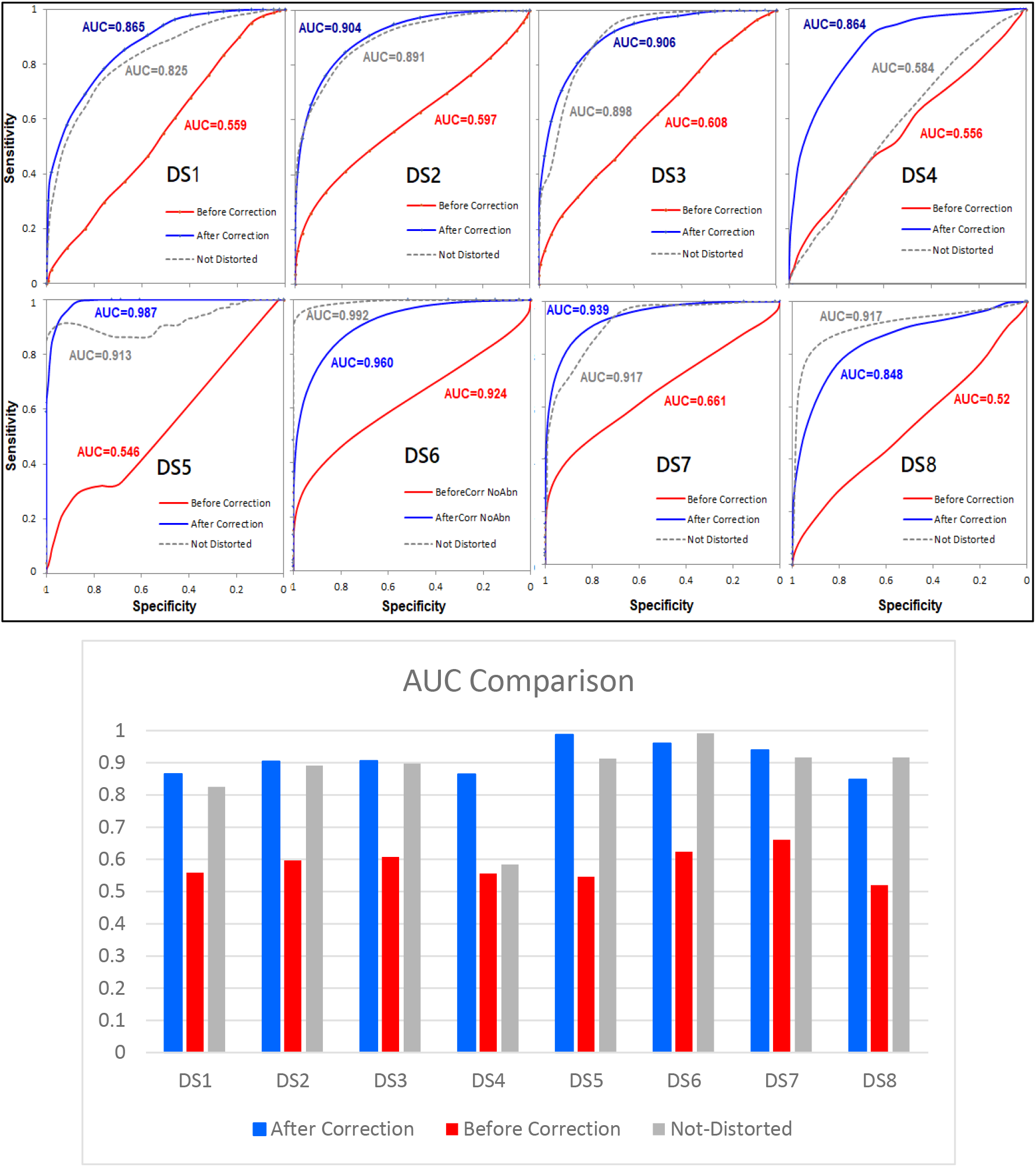
Cross Validation Results. The above plots show the prediction cross validation results on the eight datasets used in this study denoted DS1 through DS8. In each plot, the red curve shows prediction before correction, while the blue curve shows prediction after correction. The grey curve shows the prediction results of the non-distorted markers in each dataset. The bar chart shows the AUC comparisons between the three curves.

#### Cross Dataset Validation

The cross dataset validation results are shown in Error! Reference source not found.. Each plot represents the results of prediction testing on a validation dataset. The four plots are in concordance with the cross-validation results. The results depict an improved predictive ability (26% on average) of the corrected markers, represented by the improved AUC after correction as opposed to the AUC before correction. In DS9, for example, there is approximately 34% improvement in the corrected markers’ predictive ability delineated by the increase of AUC from 0.549 before correction to 0.888 after correction.

#### Comparison with other machine learning algorithms

Two state of the art predictive models, Random Forest and Support Vector Machine (SVM), were used to compare their results to the prediction results of the introduced Spotter-Marker model.

The two models were used with default optimized parameters, for a fair comparison. **Figures 12**Error! Reference source not found. shows the prediction results of the Spotter-Marker model compared to that of Random Forest and Support Vector Machine predictive models.

**Figure 11.**
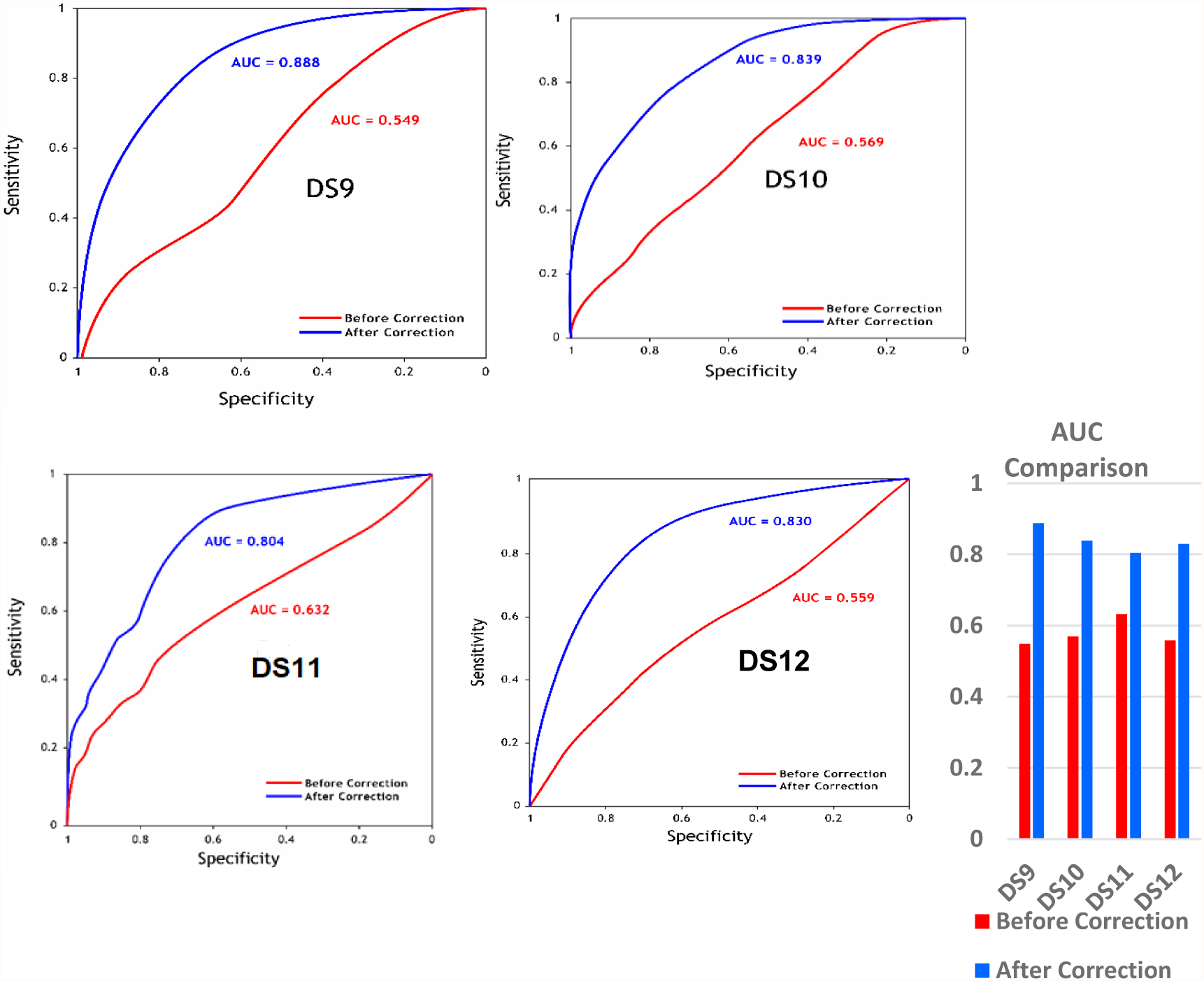
Cross dataset prediction testing. The above plots show the results of prediction testing on four independent gene expression (validation) datasets. In each plot, the red curve shows prediction before correction, while the blue curve shows prediction after correction. The four plots demonstrate the prediction improvement after correction. The bar chart on the side, compares the AUC before and after correction in each dataset.

**Figure 12.**
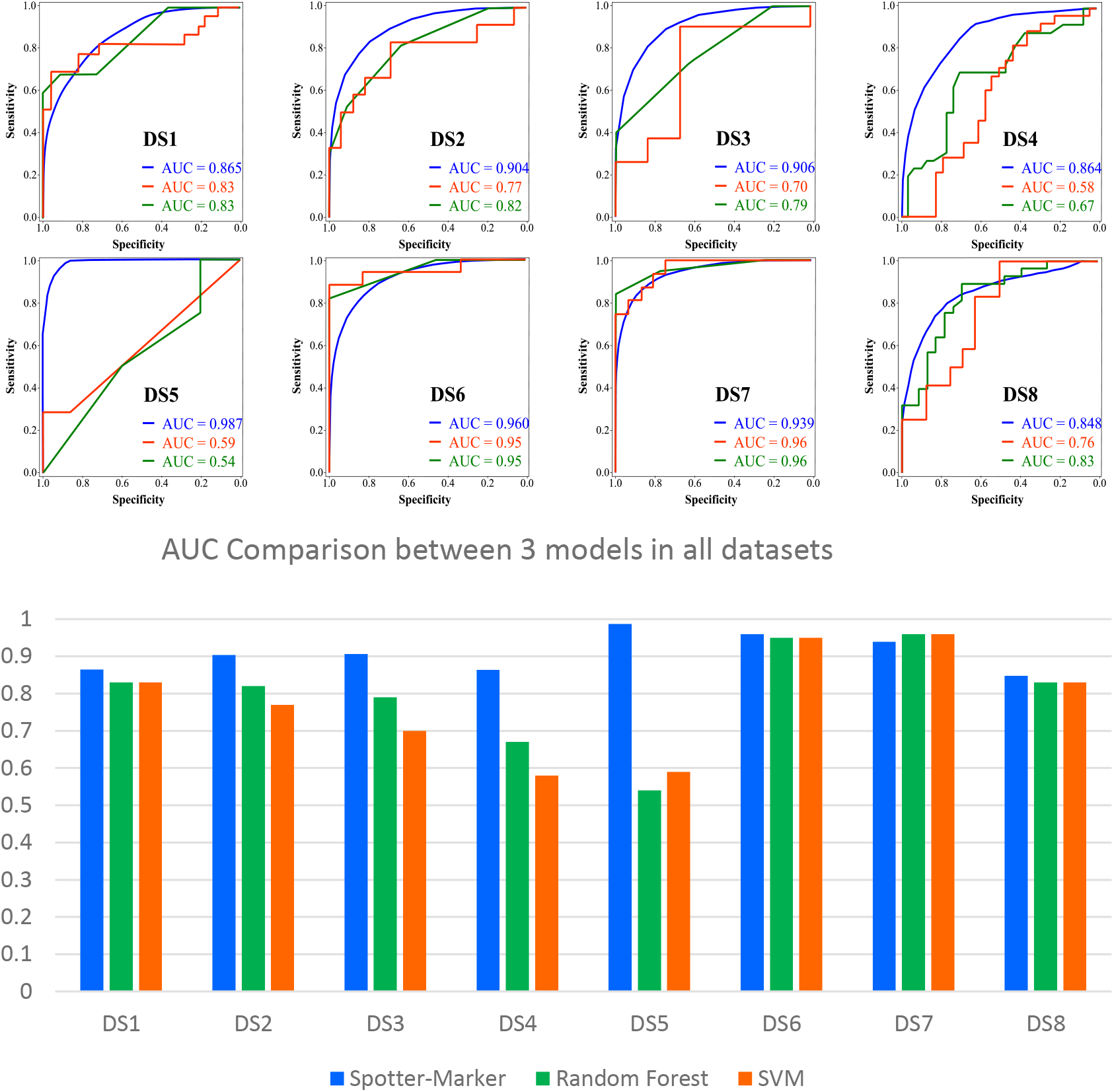
Prediction Performance: Comparison with Random Forest and SVM. The eight plots show the ROC curves of the prediction results of the Spotter-Marker model (Blue) compared to the results of Random Forest (Green) and SVM (Orange) predictive models on the eight gene expression datasets denoted DS1 through DS8. The AUC associated with each ROC curve is shown in the same color as the curve. The bar chart in the lower panel compares the AUC of each model in each dataset.

As shown in **Figure 12**, the Spotter-Marker model provides a better prediction, as demonstrated by the area under the receiver operating characteristic curves (AUC) across all datasets except DS7 in which the AUC of Spotter-Marker model is slightly less by 2%. In DS5, for example, our model shows an improvement in predictive power of 40% over that of the SVM and an improvement of 47% over that of Random Forest.

In-spite of the fact that the compared to models (Random Forest and Support Vector Machine) are popular ones in our problem domain, and that they use so many features, our model, in a much simpler fashion, achieves better results. Moreover, it adds another dimension, by explaining the biological relevance of the hidden factors behind distortion, shedding light on its sources; thus providing an interpretable predictive model.

By examining the feature importance lists of both Random Forest and SVM; that shows which features had the highest weight in predictions, to produce the above results, it was observed that the two machine learning algorithms did not use any of the discovered distorted markers. Moreover, both algorithms, used a few of the top spotters, but with much lower importance.

To test the output of the two machine learning algorithms, once in case they were not to use any of our top spotters, and another time, in case they would only use our discovered distorted markers and their spotters, we perform two tests:

1. Create eight new datasets (test 1 datasets; corresponding to the eight original datasets) which do not have any of the discovered top spotters, then train two predictive models, one using Random Forest and the other using Support Vector Machine, on the new datasets.
2. Create eight new datasets (test 2 datasets; corresponding to the eight original datasets) each contains our discovered distorted markers and their spotters only from their corresponding original dataset. Train two predictive models, one using Random Forest and the other using Support Vector Machine, on the new datasets.

The results of the above two tests are shown in **Figures 13,14**. The figures show the results of the first test using Random Forest and SVM respectively. **Figure 13** shows the results of Random Forest, and it demonstrates an improved AUC in most datasets, except DS3 which shows a slight decrease of 7% and DS7 had no change in AUC. **Figure 14** shows the results of SVM, and illustrates an improved AUC in most datasets, except DS3 and DS5 which show a decrease in AUC of 12% and 9% respectively.

**Figure 13.**
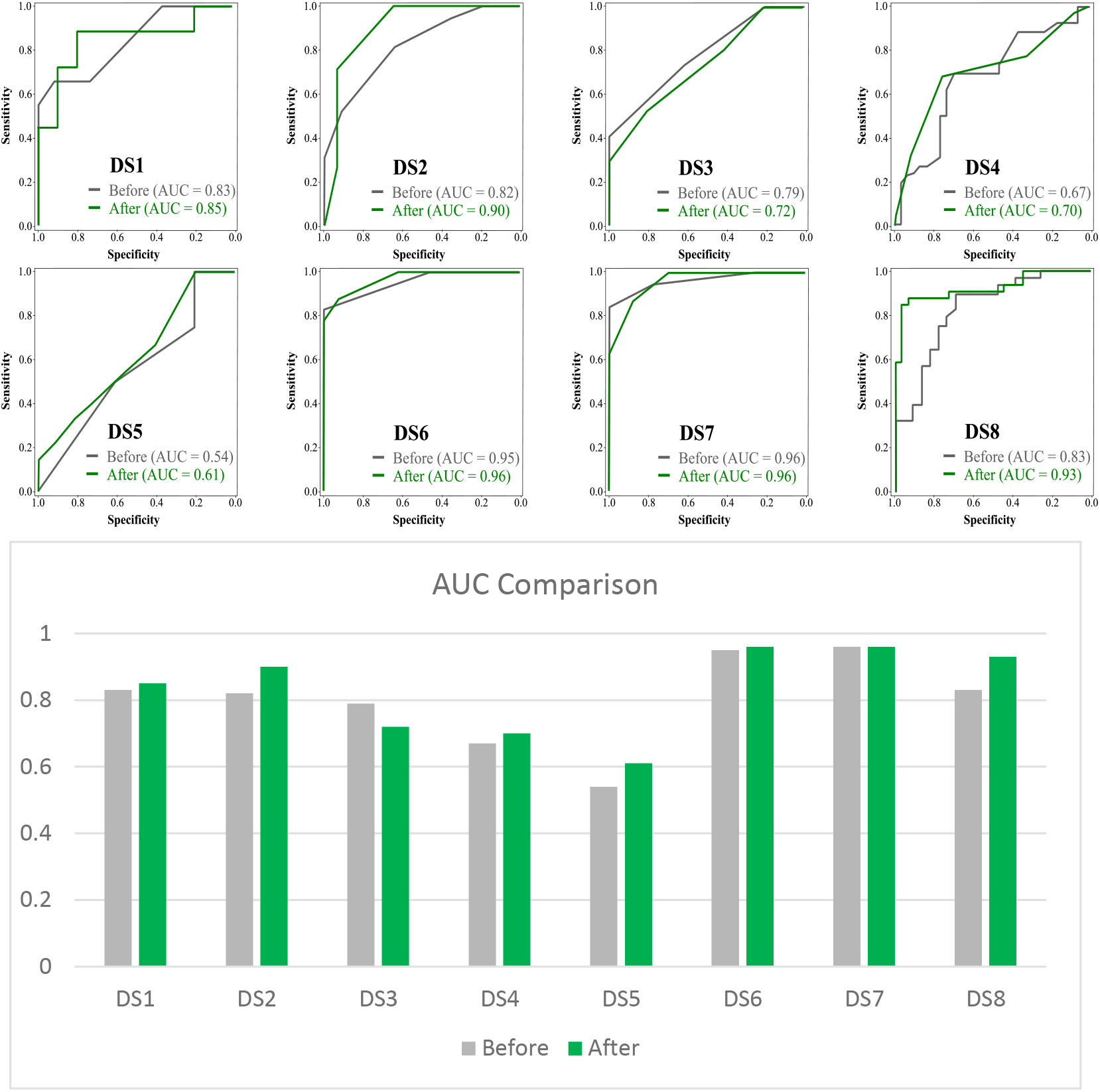
Comparison of prediction performances of Random Forest before and after removal of all top spotters from the eight datasets. The figure shows the prediction results of Random Forest on each of the eight datasets (DS1-DS8) before (Grey) and after (Green) the removal of the top spotters. The bar chart, in the lower panel, shows the AUC comparisons of (Before) and (After) across the eight datasets.

**Figure 14.**
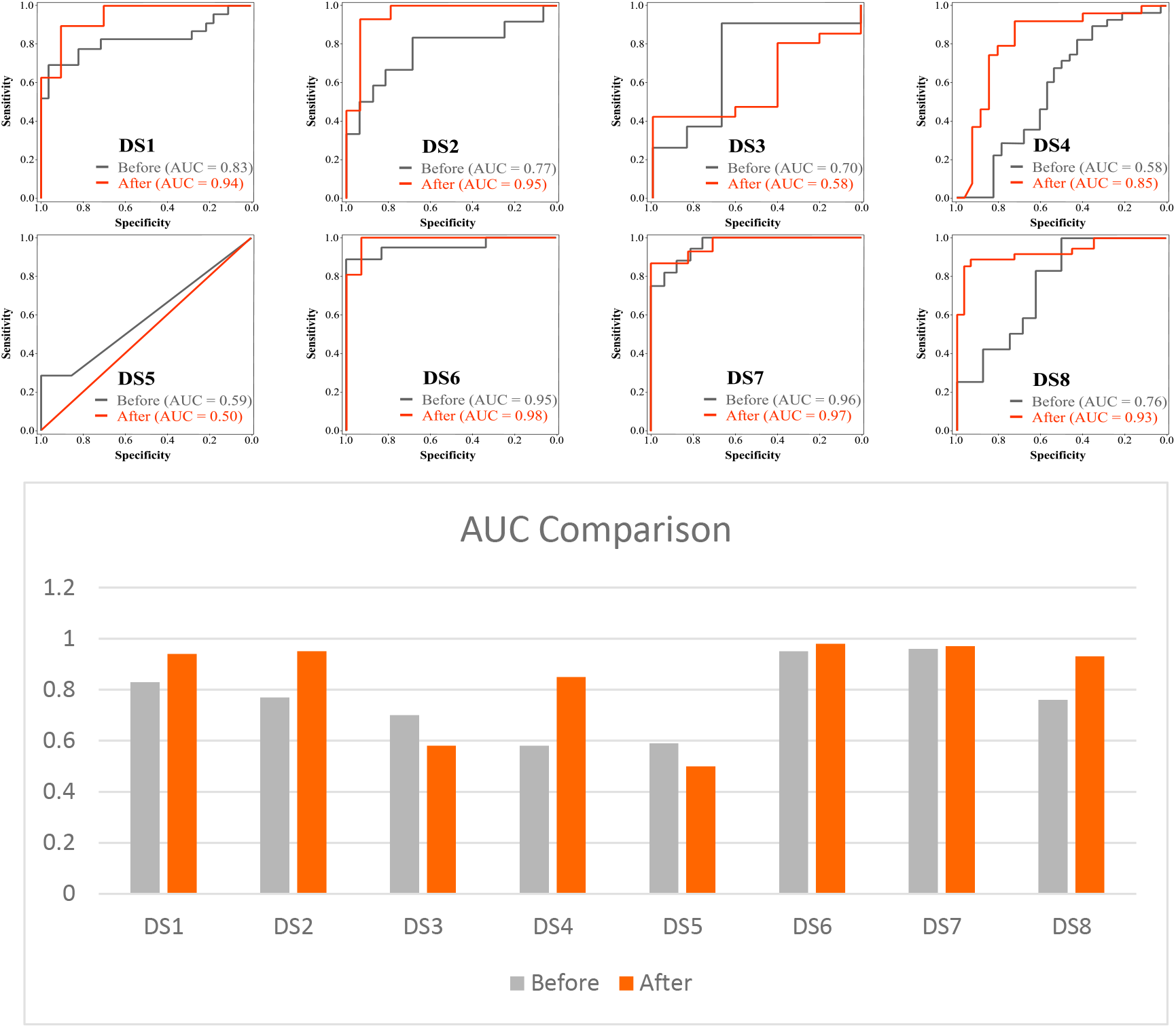
Comparison of the prediction performances of SVM before and after removal of all top spotters from the datasets. The figure shows the prediction results of SVM on each of the eight datasets (DS1-DS8) before (Grey) and after (Orange) the removal of the top spotters. The bar chart, in the lower panel, shows the AUC comparisons of (Before) and (After) across the eight datasets.

**Figures 15,16** demonstrate the results of the second test using Random Forest and SVM respectively. The two figures show consistent improvements of prediction results, demonstrated by an increased AUC in all datasets. For example, the two figures show a big improvement of 27% in AUC using Random Forest on DS5, and a bigger improvement of 31% in AUC using SVM on the same dataset. This means that the two models (Random Forest and SVM) performed better, on the new datasets that had the previously discovered spotters and distorted markers and that had none of the perfect markers that were initially selected by the two predictive models.

**Figure 15.**
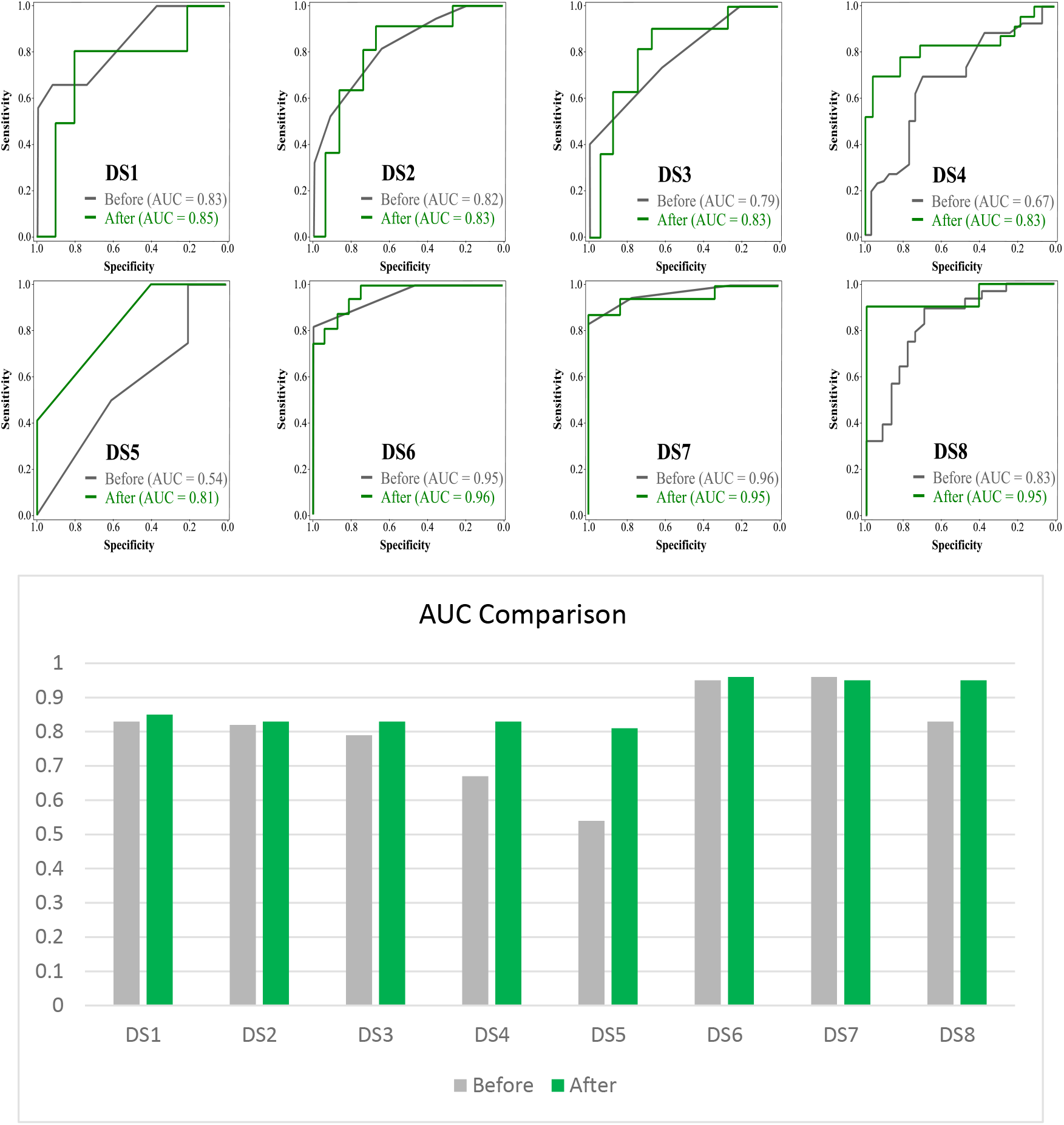
Prediction Results of Random Forest when using spotters and distorted markers as the only features in the dataset. The figure shows a comparison between the prediction results of Random Forest machine learning algorithm on all datasets (DS1-DS8), as opposed to its performance on eight new datasets created from the spotters and distorted markers discovered in each of the eight datasets respectively. Each plot shows the ROC of the prediction testing on the original dataset in grey color and the ROC resulting from prediction testing on the new dataset is shown in green color. The bar chart, in lower panel, shows the AUC comparisons of (Before) and (After) across the eight datasets.

**Figure 16.**
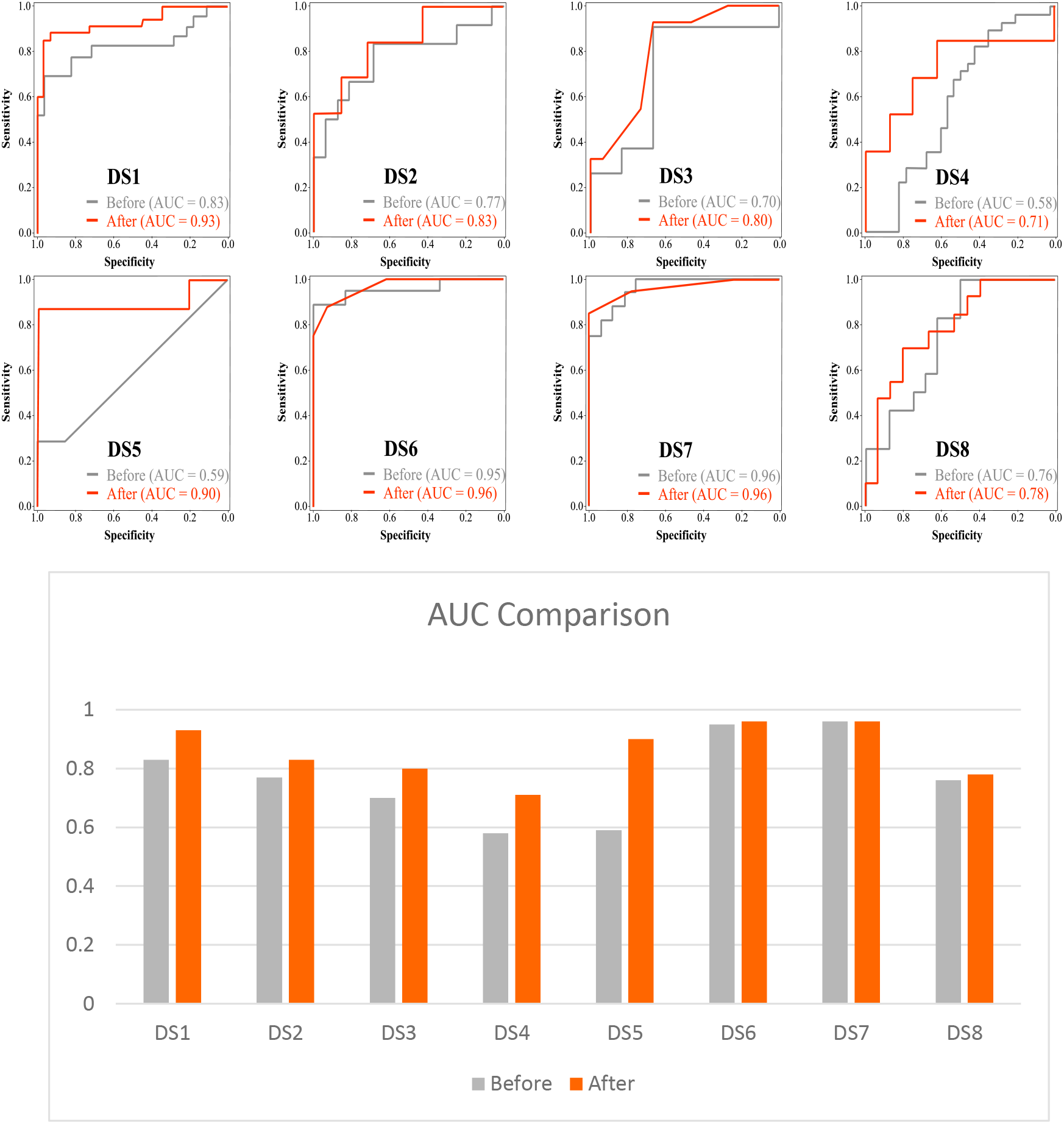
Prediction Results of Support Vector Machine when using spotters-markers as the only features in the dataset. The figure shows a comparison between the prediction results of SVM learning algorithm on all datasets (DS1-DS8), as opposed to its performance on eight new datasets created from the spotters and distorted markers discovered in each of the eight datasets respectively. Each plot shows the ROC of the prediction testing on the original dataset in grey color and the ROC resulting from prediction testing on the new dataset is shown in orange color. The bar chart shows the AUC comparisons of (Before) and (After) across the eight datasets.

### 4.3.6 Biological Relevance

#### Overlap analysis

In this section, we present the overlap across the eight gene expression datasets between: 1. The discovered spotter genes 2. The marker genes and 3. The spotter-marker pairs. For each one, we compute (i) an overlap matrix and (ii) a cumulative overlap function. Each overlap matrix is a *k*×*k* matrix that represents the pairwise overlap between the phenotype of interest -associated entity (spotter gene, marker gene, spotter-marker pair) in pairs of datasets, where *k* is the number of datasets. Each cumulative overlap function is a function in the form *f*:{1, *…, k*} → [0,1], assessing the fraction of entities (1. spotter genes 2. marker genes and 3. spotter-marker pairs) that are found to be associated with the phenotype of interest in at least a given number of the datasets. Notice that in (i) we use *k* = 8, while in (ii) we use *k* = 3 because in the latter we only asses the cumulative overlap between the three datasets of the exact same biology i.e. Prostate Tumor versus Normal.

We hierarchically cluster the datasets using each of the three overlap matrices and visualize the overlap matrices as heatmaps with hierarchical clustering. To assess the statistical significance of the overlap functions, we report the results of permutation tests obtained through 1000 permutations (the procedure we use for the permutation tests is described in Methods). We compare the overlap function computed on the original dataset against the distribution of overlap functions computed using permutation tests, representing one thousand simulated runs.

The spotter gene overlap matrix for prostate cancer in the eight datasets and the spotter gene cumulative overlap function for prostate tumor versus normal prostate tissue are shown in **Figure 17(a)** and **Figure 18(a: blue)**, respectively. As seen in the figures, the overlap between the datasets of the same biology of interest seem to be highest, i.e. between (DS1, DS2, DS8) and between (DS4, DS7). There is also some overlap between (DS2, DS6 and DS7) which are coming from the same experiment. Moreover, the permutation test for the overlap function, Error! Reference source not found.**(b)** for k=2 and k=3(two or three datasets of the same biology) suggests that the pairwise overlap is statistically significant (*p* < 0.001). In other words, a spotter gene that is found to be associated with prostate tumor versus normal prostate in one dataset is likely to be found associated with the same phenotype in at least two other datasets of the same biology.

**Figure 17.**
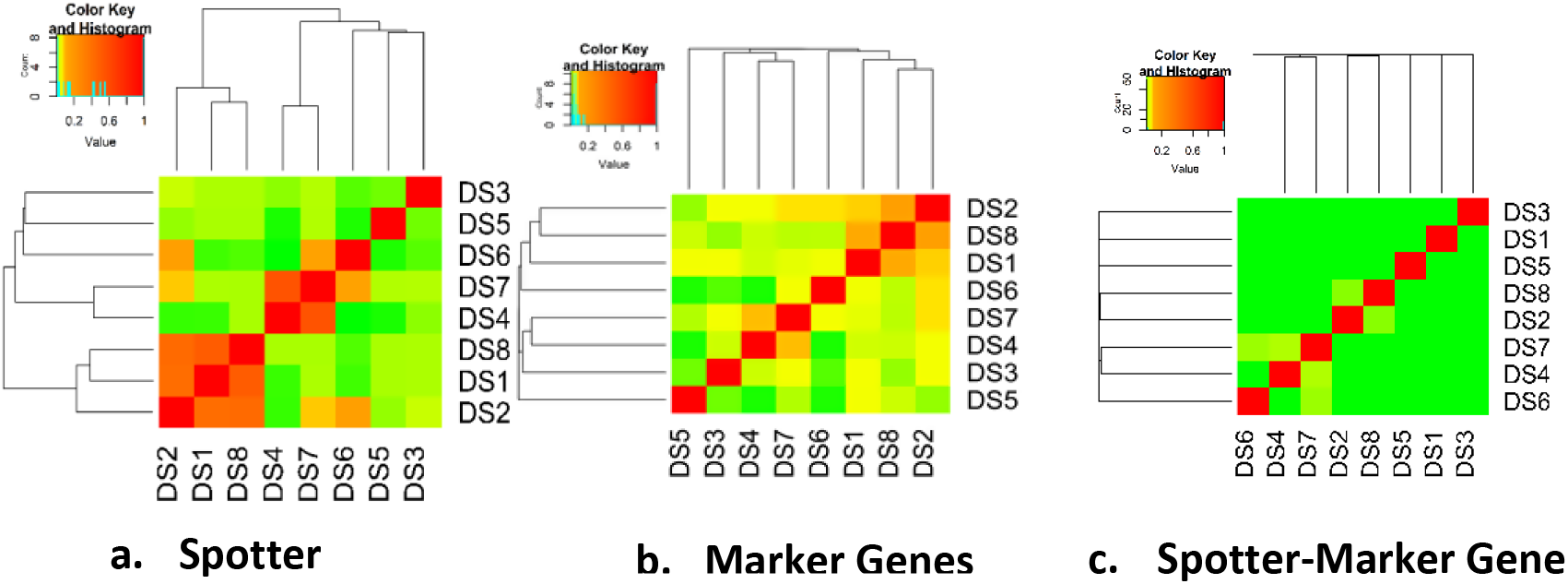
Pairwise overlap of spotter genes, marker genes and distortion instance across eight different gene expression datasets. The plots from left to right (a-c) show the overlap matrices for spotter genes, marker genes and spotter-marker instances respectively across eight datasets, depicted as heatmaps. Each heatmap has hierarchical clustering of the datasets (DSx=dataset #x). The color intensity from green to yellow to red, illustrates the degree of overlap, with red representing the highest overlap.

**Figure 18.**
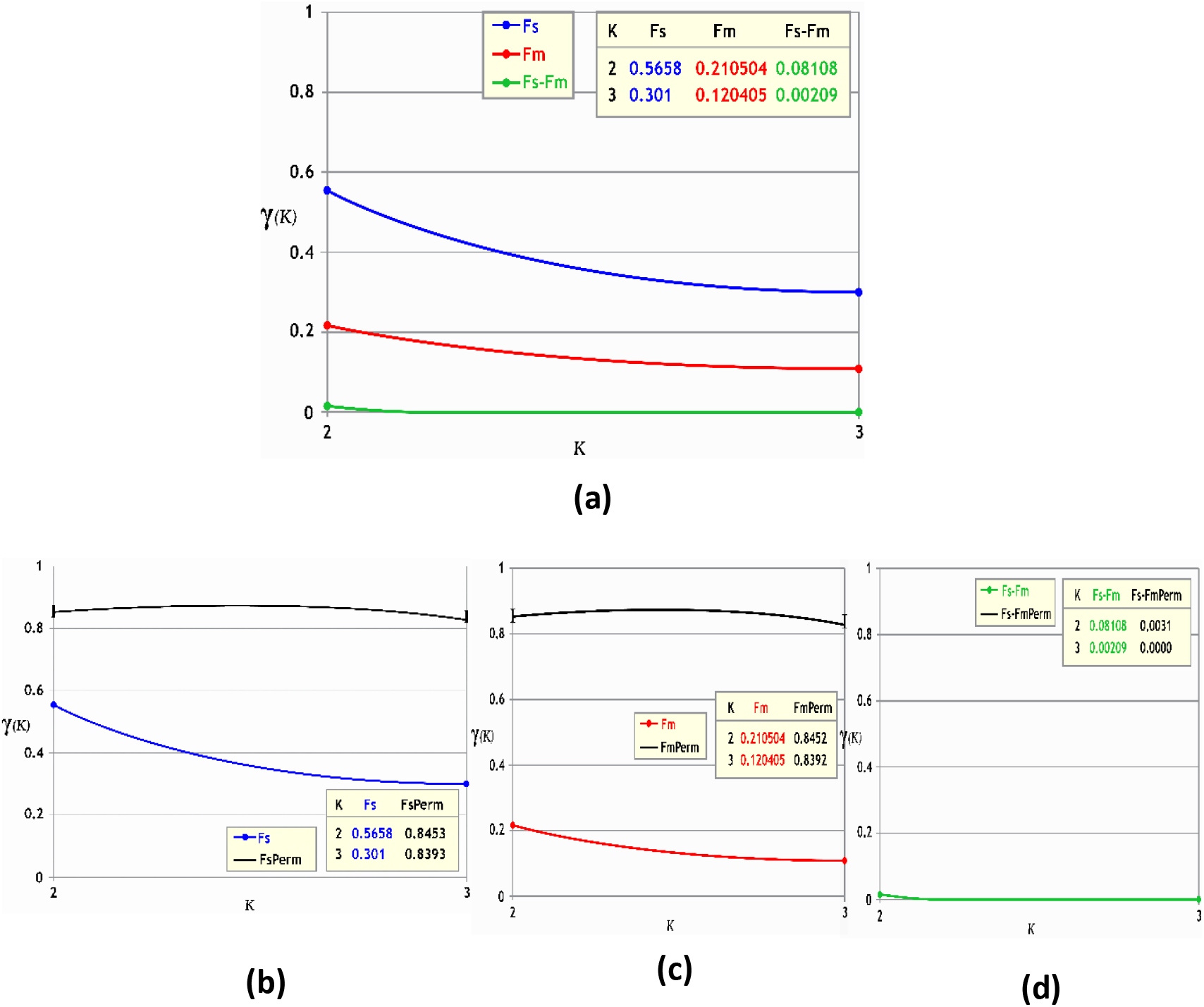
Cumulative overlap functions of spotter genes, marker genes and distortion instances across three datasets of the same phenotype. The top plot (**a**), shows the cumulative overlap functions for the spotter genes (blue), marker genes (red) and spotter-marker pairs (green) that are found to be associated with Prostate Primary Tumor versus Prostate Normal tissues in three datasets. The lower plots (**b-d**), each provides a comparison to the corresponding cumulative overlap functions computed over 1000 random permutations (black), for the spotter genes, marker genes and spotter-marker pairs, respectively.

The marker gene overlap matrix for prostate cancer in the eight datasets and the marker gene cumulative overlap function for prostate tumor versus normal prostate tissue are shown in Error! Reference source not found.**(b)** and **Figure 18 (a: red)**, respectively. As seen in the figures, the overlap between the datasets of the same biology of interest seem to be highest, i.e. between (DS1, DS2, DS8) and between (DS4, DS7). There is also some overlap between (DS2, DS6 and DS7) which are coming from the same experiment. Overall, there is more spread out pairwise overlap between prostate marker genes, across different subclasses of prostate cancer, however, overlap intensity is less focused than in the case of spotter genes overlap. This is also demonstrated by the marker gene cumulative overlap function. Moreover, the permutation test for the overlap function, **Figure 18 (c)** for k=2 and k=3(two or three datasets of the same biology) suggests that the pairwise overlap is statistically significant (*p* < 0.001). In other words, a marker gene that is found to be associated with prostate tumor versus normal prostate in one dataset is likely to be found associated with the same phenotype in at least two other datasets of the same biology.

The spotter-marker gene pairs, i.e. the distortion instance, overlap matrix for prostate cancer in the eight datasets and the spotter-marker gene pairs cumulative overlap function for prostate tumor versus normal prostate tissue are shown in Error! Reference source not found.**(c)** and **Figure 18 (a: green)**, respectively. As seen in the figures, the overlap is much less at the distortion instance level even between the datasets of the same biology of interest, i.e. between (DS1, DS2, DS8) and between (DS4, DS7). Moreover, the permutation test for the overlap function, **Figure 18**Error! Reference source not found.**(d)** for k=2 and k=3(two or three datasets of the same biology) suggests that the pairwise overlap is not statistically significant. In other words, a distortion instance (i.e. spotter-marker gene pair) that is found to be associated with prostate tumor versus normal prostate in one dataset is unlikely to be found associated with the same phenotype in another dataset of the same biology.

## 4.4 DISCUSSION

In order to understand the biological relevance of the discovered spotter genes, the significant top spotter genes that were discovered in the eight gene expression datasets were matched against the stromal and immune signatures previously defined by Yoshihara et al. ^4^

As shown in **Figure 19**, the top spotters identified in each of the prostate cancer datasets overlap with the previously defined immune and stromal signatures at varying degrees. The network of the top spotters identified in the eight datasets represent 40% overlap with the previously defined Stromal signature, 49% of the previously defined Immune signature and 49% of both signatures. The lower panel demonstrates the corresponding results computed as the average of a randomly selected set of genes equal size to the number of spotters identified from each dataset, repeated a thousand times. This finding suggests that the spotter genes may be associated with infiltrates of stromal and immune system impurities in the tumor tissue as suggested by Yoshihara et al. ^4^ hence lowering the tumor signal and resulting in higher false negative rates. **Figure 19,** also suggests that the spotter genes could be associated with some other undiscovered biological functions as well.

**Figure 19.**
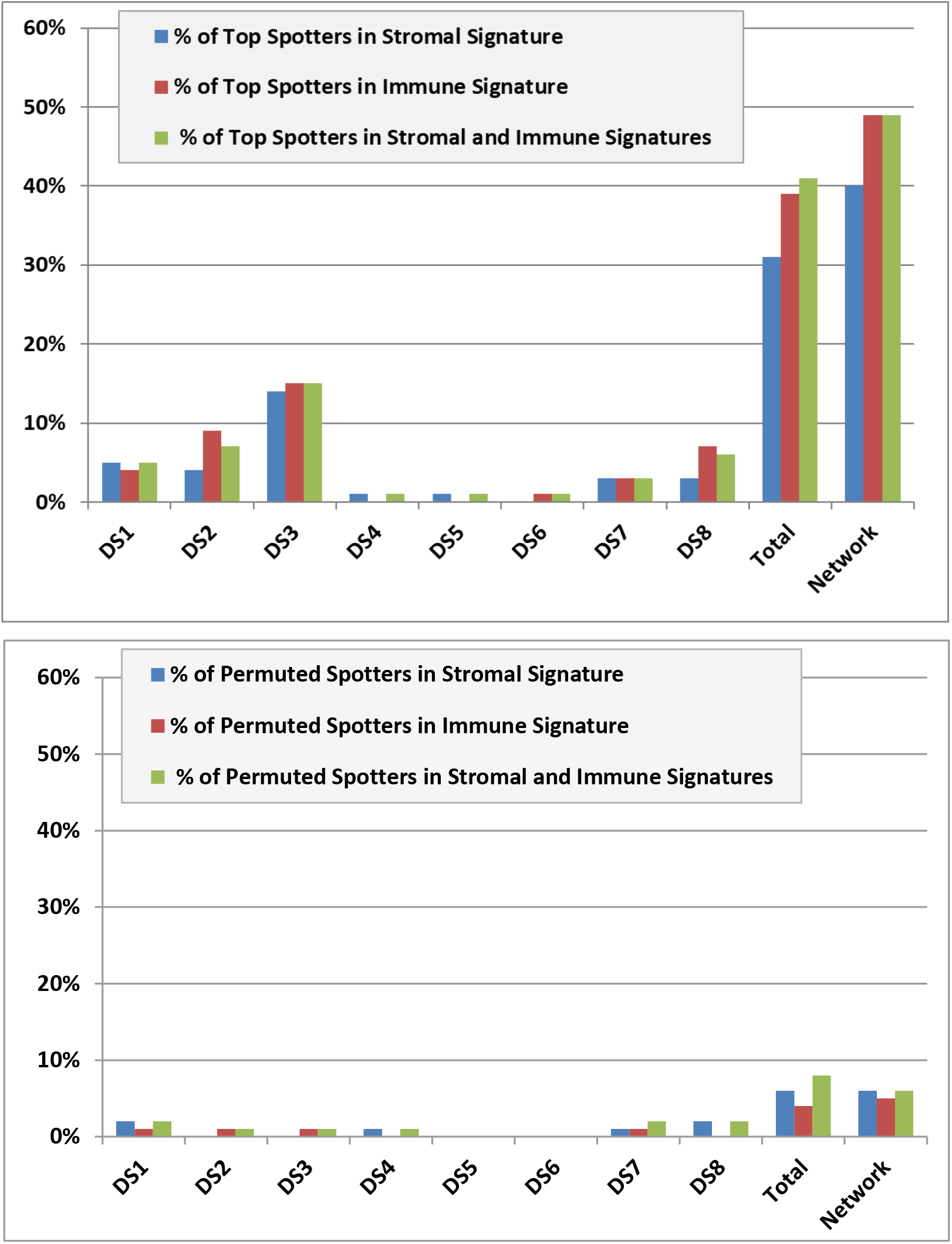
Percentage of discovered top spotter genes in Immune and Stromal signatures. The figure plots the fraction of: 1. The top spotter genes discovered on each dataset, 2. The total top spotter genes discovered in the eight datasets and 3. The genes in the network of top spotters discovered in the eight datasets, to the immune and the stromal signatures. The lower panel plots the corresponding average fractions over 1000 permuted data.

On the other hand, the results of the overlap analysis demonstrate some coherence in the biological function of those discovered spotter genes. For example, the spotter gene *MT3*, which is found in 7 datasets, is a modulator in immunity and infection. The spotter genes shared among 6 datasets, are associated with regulation of stem cell proliferation, and are related to immune cell function. The central gene in the distortion instances (F_s_-F_m_) network is *CD200* is a protein coding gene that belongs to the immunoglobulin superfamily. Moreover, the overlapping gene in the top spotters across the eight datasets used in this study, *ABCB8*, is associated with disruption in iron homeostasis. The role of iron in immunity is necessary for immune cells proliferation and maturation, particularly lymphocytes, associated with the generation of a specific response to infection ^43, 44^.

The results of in-depth analysis, where all distortion instances of each dataset are searched (and grouped) for (by) common distorted samples, indicate functional coherence among the common spotter genes that spot the same group of distorted samples within each dataset. The spotter genes that were commonly identified spotting distortion on the largest groups of distorted samples show highly coherent co-expression networks in all of the eight datasets. In some of the datasets, like DS1 for example, such networks are enriched in immune system related functionalities like regulation of purine nucleotide biosynthesis, and purine metabolism.

## Conclusion

This work introduces a new method that models the relationships between disease biomarkers and other non-marker genes; which is called Spotter genes. The introduced Spotter-Marker model improves disease risk prediction from gene expression data, by spotting, explaining and mitigating a portion of the inherent noise (typically, distortion caused by unknown factors). For this purpose, Prostate Cancer is chosen as a model complex disease. An end-to-end method is developed: from hypothesis forming to experimentations and validation of results. The results support the hypothesis with high significance, and confirms that the introduced model quantitatively and qualitatively improves disease risk prediction by unveiling new, otherwise missed biomarkers after spotting and correcting for the effect of distortion in eight prostate cancer datasets of different biology (case/control, metastatic/non-metastatic, aggressive/non-aggressive, etc.).

Correcting for the top scoring statistically significant distortion instances show significant improvements in the mutual information of the distorted markers, this is further demonstrated by the results of risk prediction testing shown in the ROC curves. High false negative rates, that are thought to be related to tumor impurity caused by the coexisting non-tumor cells, are dramatically lowered. The statistically significant distortion instances seem to have biological relevance demonstrated by its overlap across different datasets, and furthermore, they have overlapping biological functions (e.g. many spotters are associated with immune functions).

The results show that the new Spotter-Marker model successfully spots and efficiently corrects the false negative predictions. The corrected markers gain higher predictive power such that they achieve AUC values comparable to, or better than, the non-affected markers.

To that end, the comprehensive experimental results presented in this work support the utility of the Spotter-Marker model. The inclusion of the spotter(s) in addition to marker(s) in the predictive model improves its predictive power. This is further validated by comparison to some of the current, state of the art, predictive models that use multiple predictive features. The introduced model is shown to produce better results, while being much simpler. In addition, the resulting predictive models are more interpretable.

The results from this work inspire some possible future avenues in the direction of explaining distortion and improving disease risk assessment. For example, defining global markers of specific types of distortion that are expected to act as spotters in any dataset. Exploring collaborative spotters that can, each, spot small number of distorted samples which are considered noise by each individual spotter (less than MinMis), or taking the spotters and markers to a many-to-many relationship in which a network of related spotters that spots the mislabeling of a group of markers can be discovered. This can be utilized by high order machine learning algorithms for dimensionality reduction.

Finally, while integration of multi-level omics data is highly promising in the sense that it can shed light on biological pathways beyond the potential of a single omics data layer, yet it introduces challenges like more noise, integration issues, and increasing the curse of dimensionality, which is already an existing problem at each omics layer by itself. There is a need for sever, but informed, dimensionality reduction before building predictive models. The introduced Spotter-Marker model can be utilized as an informed feature selection method, that exploits knowledge of the spotter and marker relationships to select discriminative features that improve the accuracy of predictive models.

